# Beyond the Anterior Temporal Lobe: Domain-Related Degeneration of Cortical Language Network Dynamics in Semantic Dementia

**DOI:** 10.1101/2025.08.29.673019

**Authors:** Junjing Li, Lin Huang, Zhe Hu, Yaling Wang, Xiaolin Guo, Junjie Yang, Jiaxuan Liu, Runxiang Yao, Binke Yuan, Qihao Guo

**Affiliations:** Key Laboratory of Brain, Cognition and Education Sciences, Ministry of Education, China: Institute for Brain Research and Rehabilitation, South China Normal University, 510631 Guangzhou, China; Department of Gerontology, Shanghai Sixth People’s Hospital Affiliated to Shanghai Jiao Tong University School of Medicine, Shanghai, China; Philosophy and Social Science Laboratory of Reading and Development in Children and Adolescents (South China Normal University), Ministry of Education Center for Studies of Psychological Application, South China Normal University, 510631 Guangzhou, China; Center for Studies of Psychological Application, South China Normal University, 510631 Guangzhou, China

**Author notes:** Correspondence to: Qihao Guo, M.D., Department of Gerontology, Shanghai Sixth People’s Hospital Affiliated to Shanghai Jiao Tong University School of Medicine, Shanghai 200233, China Address: No.600, Yishan Road, Xuhui District, Shanghai, China Postcode: 200233, Binke Yuan, Ph.D., Institute for Brain Research and Rehabilitation, South China Normal University, Guangzhou, China Address: No.55, West of Zhongshan Avenue, Tianhe District, Guangzhou City Postcode: 510631.

## Abstract

Semantic dementia (SD) is a neurodegenerative disorder marked by a progressive decline in semantic memory, mainly due to focal atrophy in the anterior temporal lobe (ATL). As the disease advances, atrophy spreads to the perisylvian regions, accompanied by non-semantic language impairments. Despite these clinical findings, the network mechanisms behind cross-domain linguistic deficits remain poorly understood. In this study, we used our recently developed meta-networking framework of cortical language network dynamics to systematically examine domain-specific network disruptions in SD. Using resting-state functional MRI (fMRI) and comprehensive neuropsychological tests across several language domains, we analyzed data from SD patients at two timepoints: baseline (n = 42) and a 2-year follow-up (n = 24). Our findings showed that the framework successfully identified domain-specific language network degeneration. Beyond the ATL, progressive atrophy disrupted the dynamic separation of language networks involved in semantic processing, phonological processing, and speech production. These disruptions were characterized by state-specific hypo- and hyper-connectivity patterns that related to distinct language impairments. At follow-up, atrophy extended to posterior temporal and prefrontal regions, worsening network function. Importantly, the patterns of language network disruption predicted individual language deficits, providing a mechanistic link between structural degeneration, functional network changes, and clinical symptoms.

## Introduction

Semantic dementia (SD) is recognized as a clinical variant of both primary progressive aphasia (sv-PPA) and frontotemporal dementia (FTD) (Gorno-Tempini et al., 2011; Neary, 1998). Clinically, SD shows a progressive loss of cross-modal semantic knowledge, including deficits in lexical-semantic processing (e.g., word understanding and naming) (Chen et al., 2017; Ding et al., 2016; Patterson et al., 2007; Snowden et al., 2018) and non-verbal object recognition (e.g., difficulty identifying objects, faces, and voices) (Goll et al., 2012; Hodges & Patterson, 2007; Lambon Ralph, 2014; Piwnica-Worms et al., 2010). The syndrome is pathologically linked to focal atrophy in the bilateral ATL, with progressive involvement of limbic and frontoinsular regions in later stages (Ding et al., 2024; Hodges & Patterson, 2007; Landin-Romero et al., 2016; Snowden, 2001). The estimated prevalence of FTD is 10.8 per 100,000, with semantic dementia accounting for about one-third of cases (Cole et al., 2024; Coyle-Gilchrist et al., 2016). Despite its rarity, SD serves as a valuable model for studying neural substrates of semantic memory, cross-modal integration (e.g., verbal/non-verbal semantic processing), and neurodegenerative selectivity (Hodges & Patterson, 2007; Landin-Romero et al., 2016).

Early studies consistently show that SD patients initially retain core non-semantic abilities, including motor speech (e.g., fluent phonology and prosody) (Caine et al., 2009; Cousins et al., 2018; Kertesz et al., 2010), sensory perception (e.g., intact visual perception and auditory processing) (Belliard et al., 2013; Snowden et al., 2018), and executive functions (e.g., working memory) (Montembeault et al., 2016). However, these abilities are usually limited to the prodromal or early stages of the disease (Belliard et al., 2013; Ding et al., 2024). As SD progresses, patients develop complex language deficits, starting from lexical retrieval problems (e.g., anomia and fluent but empty connected speech) (Hodges et al., 1995; Mesulam et al., 2012) to widespread impairments in phonological processing (e.g., phoneme discrimination) (Bright et al., 2008), syntactic understanding (Bright et al., 2008; Heitkamp et al., 2016), and morphological rules (Heitkamp et al., 2016; Lerman et al., 2023). At the same time, cognitive decline becomes apparent, with decreases in global cognition (MMSE scores) (Huang et al., 2023), sound perception (Goll et al., 2012; Huang et al., 2023), and episodic memory (Chen et al., 2019; Ding et al., 2016). Longitudinal neuroimaging shows widespread atrophy extending beyond the ATL, including the bilateral insula, posterior temporal lobe, inferior frontal gyrus, middle frontal gyrus, medial prefrontal cortex, and posterior cingulate cortex (Hodges & Patterson, 2007; Huang et al., 2023; Landin-Romero et al., 2016). Diffusion tensor imaging-based tractography studies indicate significant reductions in the integrity of the inferior longitudinal fasciculus (ILF), arcuate fasciculus, superior longitudinal fasciculus (SLF), uncinate fasciculus, and corpus callosum (Agosta et al., 2010; Chen et al., 2020). While the semantic deficit network (Binney et al., 2010; Patterson et al., 2007; Wilson et al., 2014) is well documented, the neural bases of non-semantic linguistic impairments (e.g., repetition/phonology) remain less studied.

Language is a multidimensional system made up of distinct, yet interconnected components, including phonetic perception, phonological representation, semantic representation, speech articulation, morphological processing, and syntactic processing (Hickok, 2022; Hodgson et al., 2021; Matchin & Hickok, 2020; Schaeffer et al., 2023). The neurobiology of language involves a distributed network across the bilateral temporal, frontal, and parietal lobes (Fedorenko & Thompson-Schill, 2014; Lipkin et al., 2022; Lu et al., 2021; Skeide & Friederici, 2016), which are connected by underlying white matter tracts (Duffau et al., 2014; Saur et al., 2008; Yeh, 2022). Although the brain regions supporting each language domain are spatially distinct, they work together as a coordinated language network organized through specific functional pathways. Both traditional language models and the dual-stream model emphasize the core components of the dorsal phonological pathway and the ventral semantic pathway (Duffau et al., 2014; Hickok, 2022). Recently, we proposed the "dynamic meta-networking framework of language" (Yuan, Xie, Wang, et al., 2023), which expands on the dynamic reorganization of the language network during resting state. This framework divides the language network into three pathways: the dorsal speech articulation pathway, the central phonological perception and representation pathway, and the ventral semantic pathway. Unlike the dual-stream model, the "dynamic meta-networking framework of language" further divides the dorsal phonological pathway into two distinct components, thereby allowing for more flexible integration within the network. Our initial research shows that the "dynamic meta-networking framework of language" is both highly reproducible and capable of predicting the extent of language impairment in patients (Yuan, Xie, Gong, et al., 2023).

In this study, we adopt the "dynamic meta-networking framework of language" to investigate domain-related linguistic degeneration in SD patients. Neuropsychological tests were administered at baseline (n = 42) and follow-up (∼2 years, n = 24), and resting-state fMRI data were collected. The linguistic batteries covered multiple domains, including cross-modal semantic processing, phonological processing, and speech production.

## Participants and Methods

### Participants

Neuropsychological and imaging data were collected from patients with SD (SDs) and healthy controls (HCs) at Huashan Hospital in Shanghai, China. All participants were right-handed, native Chinese speakers who provided written informed consent. This study was approved by the Ethics Committees of Huashan Hospital (Approval number: 2009-195) and Shanghai Sixth People’s Hospital (Approval number: 2022-ky-116). Diagnoses were made through consensus using the current diagnostic criteria (Gorno-Tempini et al., 2011), following comprehensive assessments including medical history, neuropsychological testing, brain MRI, and 18F-FDG PET scans. Overall cognitive function was evaluated with the Chinese version of the Mini-Mental State Examination (MMSE) (Katzman et al., 1988).

A total of 42 patients (24 women, 18 men; age: mean ± SD = 62.55 ± 7.11 years; MMSE: mean ± SD = 21.97 ± 4 scores) were included in this study. All patients showed atrophy in the ATL with no history of stroke, head trauma, substance abuse, severe visuo-perceptual impairment, psychiatric, or other neurological diseases. Twenty-four of these patients, who visited the memory clinic at Huashan Hospital in Shanghai, China, from 2011 to 2018, were recruited for a 2-year follow-up (interval months: 22.83 ± 10.63). A control group of 37 cognitively normal participants (18 women; average age: 61 ± 4.39 years) was selected. They performed normally on the MMSE (average score: 28.29 ± 1.36).

### Neuropsychological assessment and preprocessing

SDs and HCs underwent identical neuropsychological assessments (Table 1) covering general mental status, episodic memory, visuospatial skills, and executive functions, along with behavioral tasks (Table 2) that involved general semantic processing, basic perception, phonological processing, and speech production. These behavioral tasks were chosen to systematically examine different parts of the language system—semantic processing, phonological processing, and speech production—while including simple perceptual tasks to control for low-level sensory influences. This setup allowed for the extraction of underlying functional components through principal component analysis (PCA), laying the foundation for later mapping of behavioral profiles to neural states.

**Table 1.** Demographics and neuropsychological tests.

| Index | HC |  | SD |  | Paired SD (baseline) |  | Paired SD (follow-up) |  | HC & SD | HC & SD (follow-up) | SD (baseline) & SD (follow-up) |
| --- | --- | --- | --- | --- | --- | --- | --- | --- | --- | --- | --- |
|  | Mean (SD) | n | Mean (SD) | n | Mean (SD) | n | Mean (SD) | n | *P-value* | *P-value* | *P-value* |
| Demographics |  |  |  |  |  |  |  |  |  |  |  |
| Age, years | 61.00(4.39) | 37 | 62.55(7.11) | 42 | 61.46(5.85) | 24 | 63.25(5.92) | 24 | 0.243 | 0.094 | NA |
| Sex (F:M) | 18:14 |  | 24:18 |  | 16:8 |  | 16:8 |  | 0.650 | 0.608 | NA |
| Education, years | 10.53(3.09) | 17 | 11.43(2.74) | 42 | 11.67(2.82) | 24 | 11.79(3.18) | 24 | 0.276 | 0.212 | NA |
| Neuropsychological tests |  |  |  |  |  |  |  |  |  |  |  |
| General mental status |  |  |  |  |  |  |  |  |  |  |  |
| MMSE (max = 30) | 28.29(1.36) | 17 | 21.97(4.00) | 31 | 22.46(3.81) | 24 | 17.17(5.06) | 24 | <0.001 | <0.001 | <0.001 |
| Language |  |  |  |  |  |  |  |  |  |  |  |
| BNT (max = 30) | 22.18(3.09) | 17 | 6.87(4.13) | 31 | 7.04(4.11) | 24 | 3.29(3.78) | 24 | <0.001 | <0.001 | 0.001 |
| Episodic memory |  |  |  |  |  |  |  |  |  |  |  |
| REY-O recall (max = 36) | 17.24(6.13) | 17 | 10.81(5.15) | 31 | 11.25(5.20) | 24 | 7.79(5.58) | 24 | <0.001 | <0.001 | 0.013 |
| Visuo-spatial function |  |  |  |  |  |  |  |  |  |  |  |
| REY-O copy (max = 36) | 34.18(2.19) | 17 | 32.61(3.57) | 31 | 32.25(3.84) | 24 | 31.58(6.18) | 24 | 0.107 | 0.107 | 0.372 |
| Executive function |  |  |  |  |  |  |  |  |  |  |  |
| STT-B, seconds | 151.88(45.87) | 17 | 181.23(72.32) | 31 | 168.76(73.89) | 21 | 193.05(81.27) | 21 | 0.138 | 0.058 | 0.220 |
| Calculation (max = 7) | 6.41(0.71) | 17 | 6.45(0.85) | 31 | 6.65(0.49) | 23 | 6.43(0.59) | 23 | 0.870 | 0.912 | 0.096 |

Abbreviations: MMSE, mini-mental state examination; BNT, Boston naming test; REY-O, Rey-Osterrieth complex figure test; STT-B, the shape trail test-part B; NA, not applicable.

**Table 2.** Baseline performance and annual decline on the primary behavioral tasks.

| Index | HC |  | SD |  | Paired SD<br>(baseline) |  | Paired SD<br>(follow-up) |  | HC & SD | HC<br>&<br>SD (follow-up) | SD (baseline)<br>&<br>SD (follow-up) |
| --- | --- | --- | --- | --- | --- | --- | --- | --- | --- | --- | --- |
|  | Mean (SD) | n | Mean (SD) | n | Mean (SD) | n | Mean (SD) | n | *P-value* | *P-value* | *P-value* |
| General Semantic task |  |  |  |  |  |  |  |  |  |  |  |
| Oral picture naming (max = 140) | 126.53(7.20) | 17 | 49.70(28.07) | 30 | 48.88(27.22) | 24 | 22.13(25.16) | 24 | <0.001 | <0.001 | <0.001 |
| Oral sound naming (max = 36) | 27.12(4.27) | 17 | 8.93(5.14) | 30 | 8.83(4.55) | 24 | 3.67(4.89) | 24 | <0.001 | <0.001 | <0.001 |
| Picture associative matching (max = 70) | 66.53(2.27) | 17 | 51.53(6.94) | 30 | 52.17(7.25) | 24 | 44.92(7.85) | 24 | <0.001 | <0.001 | <0.001 |
| Word associative matching (max = 70) | 67.18(1.59) | 17 | 53.13(8.28) | 30 | 53.13(8.67) | 24 | 45.54(8.02) | 24 | <0.001 | <0.001 | <0.001 |
| Word-picture verification (max = 70) | 67.47(1.81) | 17 | 43.73(16.00) | 30 | 44.38<br>(16.38) | 24 | 28.25 (14.82) | 24 | <0.001 | <0.001 | <0.001 |
| Naming to definition (max = 70) | 58.53(6.10) | 17 | 18.40(13.78) | 30 | 17.96(14.35) | 24 | 6.42(8.89) | 24 | <0.001 | <0.001 | <0.001 |
| Elementary Perception task |  |  |  |  |  |  |  |  |  |  |  |
| Sound perception (max = 44) | 38.94(4.21) | 17 | 35.65(4.96) | 31 | 36.63(4.33) | 24 | 34.88(5.06) | 24 | 0.025 | 0.010 | 0.175 |
| Visual perception (max = 30) | 27.06(1.68) | 17 | 27.90(2.51) | 31 | 28.04(1.46) | 24 | 27.42(1.56) | 24 | 0.221 | 0.487 | 0.065 |
| Phonological Processing task |  |  |  |  |  |  |  |  |  |  |  |
| Auditory lexical decision (max = 12) | 9.63(0.72) | 16 | 7.86(2.03) | 29 | 8.10(2.56) | 10 | 6.40(2.84) | 10 | <0.001 | 0.006 | 0.242 |
| Speech repetition task |  |  |  |  |  |  |  |  |  |  |  |
| Repetition(max=12) | 11.76(0.44) | 17 | 11.34(0.86) | 29 | 11.80(0.42) | 10 | 11.20 (0.63) | 10 | 0.033 | 0.011 | 0.051 |

Each task was conducted in separate sessions on a PC using the DMDX program (Forster & Forster, 2003). These tasks have been employed in our recent studies (Ding et al., 2016; Han et al., 2013; Y. Zhao et al., 2017). Participants were tested individually in a quiet room. Each session lasted no more than 2 hours, with rest breaks available upon request.

#### General mental status

The Chinese version of the MMSE (Katzman et al., 1988) was used to assess overall mental function.

#### Episodic memory and visuospatial function

Episodic memory was assessed using the Rey-Osterrieth Complex Figure Recall test (Osterrieth, 1944; Rey, 1941), while visuospatial function was evaluated with the Rey-Osterrieth Complex Figure Copy test (Osterrieth, 1944; Rey, 1941), respectively.

#### Executive function

Executive function was assessed using the Shape Trail Test-Part B (STT-B; Q. Zhao et al., 2013) to measure cognitive flexibility, recorded by completion time (in seconds), along with a calculation task consisting of seven arithmetic problems (two addition, two subtraction, two multiplication, one division).

#### General semantic processing

Detailed tests across multiple language domains were conducted for each patient. To evaluate semantic memory, participants completed six general semantic tasks: oral picture naming, oral sound naming, picture associative matching, word associative matching, word-picture verification, and naming to definition. These tasks measured semantic knowledge through various input and output modalities (see Supplementary Material for details).

Elementary perception was evaluated using computerized visual and auditory perception tasks. Since these tasks did not involve language processing, they served as a domain-general reference, allowing us to compare language-specific impairments with intact or unrelated perceptual abilities in subsequent brain–behavior analyses.

Phonological processing was evaluated through an auditory lexical decision task, in which participants determined whether each of 12 spoken items was a real word or a pseudoword.

#### Speech repetition

Speech production was assessed using an oral repetition task that required verbatim reproduction of auditorily presented stimuli, including eight monosyllabic words and four syntactically simple sentences.

Given demographic differences such as age, sex, and education, raw scores were standardized to account for these variables using a regression-based approach (Crawford & Garthwaite, 2006). A linear regression model was created from the healthy control group data to predict expected scores for each task based on demographic variables. Discrepancy scores for SDs were then determined as the difference between observed and predicted scores. Standard error adjustments were applied to calculate standardized t-scores. This method allowed for the correction of demographic influences, providing a clearer measure of each patient’s impairment.

### Task performance through data reduction with PCA

To identify underlying cognitive components, PCA was performed on t-scores from the nine cognitive tasks using SPSS 27. The appropriateness of the data for factor analysis was confirmed through the Kaiser-Meyer-Olkin (KMO) measure and Bartlett’s Test of Sphericity. Components with eigenvalues greater than 1 were retained, and maximum variance rotation was applied to improve interpretability.

### Imaging Data Collection and MRI Settings

All participants were scanned using a 3T Siemens MRI scanner at Huashan Hospital. The data collected included an anatomical T1 scan, a diffusion-weighted scan, and a resting-state functional scan. All MRI scans were performed with the same scanner, including follow-up scans. The parameters for the structural scans were as follows: MPRAGE sequence, sagittal plane, repetition time = 2300 ms, echo time = 2.98 ms, flip angle = 9°, matrix size = 240 × 256, field of view = 240 mm × 256 mm, 192 slices, and voxel size = 1 mm × 1 mm × 1 mm. For blood oxygenation level-dependent (BOLD) resting-state functional MRI, the parameters were as follows: EPI sequence, transverse plane, repetition time = 2000 ms, echo time = 35 ms, flip angle = 90°, matrix size = 64 × 64, field of view = 256 mm × 256 mm, 33 slices, slice thickness = 4 mm, voxel size = 4 mm × 4 mm × 4 mm, with a total scan time of 400 seconds. During the scans, participants were instructed to lie still, stay awake with their eyes closed, and avoid systematic thinking.

### Imaging Data Preprocessing

The resting-state fMRI data were preprocessed through the following steps: 1) delete the first 10 volumes; 2) slice-timing correction was applied < 3 mm or 3; 3) perform head motion correction (< 3 mm or 3 degrees); 4) coregister images; 5) segment T1 images and normalize into the MNI space using DARTEL; 6) normalize functional images using the deformation field of T1 images; 7) smooth data with a Gaussian kernel (full-width at half-maximum = 6 mm); 8) apply linear detrend; 9) regress out nuisance signals (24 motion parameters, including x, y, z translations and rotations (6 parameters), their derivatives (6 parameters), quadratic terms of 12 parameters, and white matter, cerebrospinal fluid, and global mean time courses); 10) perform temporal band-pass filtering (0.01–0.1 Hz).

To illustrate the brain atrophy in the patients, T1 images were first segmented into grey matter, white matter, and CSF with a 1.5mm × 1.5mm × 1.5mm resolution. The output images were then normalized into MNI space using the DARTEL registration method. The grey matter images were further modulated and smoothed with an 8-mm full-width at half-maximum Gaussian kernel to obtain grey matter volume images.

Functional and structural imaging processing was implemented by using DPABI (https://github.com/Chaogan-Yan/DPABI) (Yan et al., 2016).

### Absolute and relative atrophy

To illustrate the absolute atrophy of grey matter volume between SDs at baseline or follow-up and HCs, two independent-sample t-tests were performed. To demonstrate the relative atrophy of grey matter volume between SDs at baseline and follow-up, a paired t-test was conducted.

### The cortical language network definition

To examine the dynamics of the cortical language network, we identified a putative cortical language network based on the Human Brainnetome Atlas (Fan et al., 2016). We extracted all parcels showing significant activation in language-related behavioral domains (i.e., language or speech) or paradigm categories (e.g., semantic, word generation, reading, or comprehension). A total of 33 cortical parcels in the left hemisphere and fifteen in the right hemisphere were selected. These regions closely match those identified in previous studies (Lipkin et al., 2022; Vigneau et al., 2006). Given the bilateral nature of language processing (Vigneau et al., 2006; Siegel et al., 2016; Turker et al., 2023; Hodgson et al., 2021) and the compensatory recruitment of right-hemisphere regions during recovery, we also included nineteen homologous parcels in the right hemisphere and one in the left hemisphere. Altogether, we defined a bilaterally symmetric cortical language network (BNL68, Supplementary Figure 1) comprising 68 cortical ROIs (34 in each hemisphere) (Yuan et al., 2022). These encompass the superior, middle, and inferior frontal gyri (SFG, MFG, and IFG), the ventral parts of the precentral gyrus (PrG) and postcentral gyrus (PoG), the superior, middle, and inferior temporal gyri (STG, MTG, and ITG), the fusiform gyrus (FuG), the para-hippocampal gyrus (PhG), and the posterior superior temporal sulcus (pSTS). The coordinates (in Montreal Neurologic Institute space) and meta-analysis results for each parcel are summarized in Supplementary Table 1.

### Framewise construction of time-varying networks using DCC

To identify the temporal reoccurring states of the language networks, the dynamic conditional correlation (DCC) approach was used to build the framewise time-varying language network (Lindquist et al., 2014). The calculation of DCC and the visualization of the results were performed using the NaDyNet toolbox (Yang et al., 2025) available at [https://github.com/yuanbinke/Naturalistic-Dynamic-Network-Toolbox].

DCC is a variant of the multivariate GARCH (generalized autoregressive conditional heteroscedasticity) model (Lebo & Box-Steffensmeier, 2008), which has proven to be particularly effective for estimating both time-varying variances and correlations. GARCH models describe the conditional variance of a single time series at time *t* as a linear combination of past values of the conditional variance and the squared process itself. All the parameters of DCC are estimated using quasi-maximum likelihood methods and do not require ad hoc parameter settings.

The DCC algorithm involves two steps. To illustrate, assume a pair of time series from two ROIs, *x*_*t*_ and *y*_*t*_. In the first step, standardized residuals of each time series are estimated using a univariate GARCH (1,1) process. In the second step, an exponentially weighted moving average (EWMA) window is applied to the standardized residuals to compute a non-normalized version of the time-varying correlation matrix between *x*_*t*_and *y*_*t*_. The mathematical expressions of the GARCH (1,1) model, DCC model, and EWMA, along with the model parameters, were provided by Lindquist et al. (Lindquist et al., 2014).

K-means clustering was adopted to decompose the dynamic functional connectivity (dFC) matrices into several reoccurring connectivity states. The best number of clusters, *k*, was determined using the elbow criterion, which looks at the ratio of within-cluster distances to between-cluster distances (Allen et al., 2014). The L1 distance function, also known as ’Manhattan distance,’ was used to measure the distance from each point to the centroid. Each time window was then assigned to one of these connectivity states.

### Topological properties of dFC states

A state-specific subject connectivity matrix was created by calculating the median of all connectivity matrices that were assigned to the same state label (Damaraju et al., 2014). Before this calculation, a correlation threshold (r > 0.2) was applied to remove weak connections that could have resulted from noise. Because the nature of negative connections is debatable (Murphy & Fox, 2017), only positive connections were included in this study. Here, we used a weighted network instead of a binarized one to preserve all connectivity information.

*<u>Global topological metrics</u>*: for each state-specific matrix, three network topological properties were calculated: 1) Total connectivity strength. The FC strength of a network was calculated by summing the functional connectivity strengths of all suprathreshold connections into one value (Nelson et al., 2017); 2) Global efficiency (gE). The gE reflects the capability for parallel information transfer and functional integration. It is defined as the average of the inverses of all weighted shortest path lengths (the minimal number of edges that one must traverse to reach another) in the thresholded matrix (Rubinov & Sporns, 2010); 3) Local efficiency (lE). The lE reflects the relative functional segregation and is defined as the average of the global efficiency of each node’s neighborhood sub-graph.

*Nodal topological metrics*: for each node, nodal strength, gE, and lE were calculated.

### Functional significance of state-dependent hub distributions

To evaluate the functional relevance of hubs in each state, we performed term-based meta-analyses using the platform NeuroQuery (https://neuroquery.org/) (Dockès et al., 2020). NeuroQuery estimates the spatial distribution of activity from 418,772 activation locations associated with 7647 terms extracted from 13,459 full-text neuroimaging articles. Five language- and speech-related terms— speech perception, inner speech, phonological processing, speech production, and semantic processing—were chosen to create meta-analytic activation maps. Each map was thresholded at Z = 3, following NeuroQuery’s standard practice, and only positive activation results were included. We assessed functional relevance by calculating Dice coefficients between binary images of hub nodes and meta-analytic results (Dice = 2* (hub * meta)/ (hub + meta)). Due to the lateralized nature of language processing, Dice coefficients were computed at both the whole-brain level and specifically within the left hemisphere.

### Statistical analysis

Age and education were compared using two-sample t-tests, and sex was assessed with Pearson’s chi-square test, all conducted in IBM SPSS Statistics version 27. Neuropsychological tests and cognitive tasks were compared between HCs and SDs at baseline using two-sample t-tests, and within SDs across baseline and follow-up using paired t-tests.

For both types of T-results maps of atrophy, FDR correction at *q* < 0.01 was applied for multiple comparisons.

The changes in functional connectivity and nodal and global topological properties were analyzed using two-sample t-tests. The edge strength results were corrected with network-based statistics (NBS) (Zalesky et al., 2010) at an edge *p*-value of 0.01 and a component *p*-value of 0.01. Nodal and global topological properties results were corrected using FDR correction at *q* < 0.05.

The associations between edge, nodal, and global topological properties and the patients’ cognitive scores were evaluated by calculating the partial Pearson correlation coefficient while controlling for age, sex, and head motion. The findings relating edge strength to cognitive scores were corrected using NBS with a component *p*-value of 0.05.

### Machine learning-based dFC-linguistics prediction model

To investigate whether the reorganized dFCs were associated with patients’ language deficits, we built multivariate machine learning-based dFC-linguistics prediction models. The relevance vector regression (RVR) algorithm and linear kernel function were adopted (Cui & Gong, 2018; Yuan et al., 2019). RVR has no algorithm-specific parameter and thus does not require extra computational resources to estimate the optimal algorithm-specific parameters (Tipping, 2001). Model performance was evaluated using leave-one-out cross-validation (LOOCV), and predictive accuracy was measured by the Pearson correlation coefficient between predicted and observed behavior scores. The statistical significance of the model was tested with 1000 permutations. Details of the RVR algorithm are provided in Supplementary Methods.

## Results

### Demographics and Cognitive Performance

Table 1 summarizes the demographics of SDs and their performance on neuropsychological tests. There are no differences in age, sex, or education between SDs and HCs. The participants’ raw scores on the behavioral tasks are shown in Table 2. After controlling for age, sex, and education, the SDs showed significant deficits in six general semantic tasks compared to HCs at baseline (*p* < 0.001). Longitudinal analyses revealed that SDs performed significantly worse at 2-year follow-up than at baseline on these semantic tasks. In addition to semantic decline, SDs also showed significantly impaired performance on sound perception and auditory lexical decision tasks compared to HCs at baseline. This suggests widespread disruptions affecting phonological processing and auditory perception, indicating broader dysfunction within the linguistic network.

### PCA of behavioral performance

All behavioral data were entered into a PCA with varimax rotation. The KMO value was 0.643, indicating moderate sampling adequacy. Bartlett’s Test of Sphericity was highly significant (p < 0.001), supporting the appropriateness of the data for dimensionality reduction through PCA. Figure 1 (A) shows the loadings of the tasks on each principal component. Three components with eigenvalues greater than 1 were extracted, explaining 77.39% of the total variance (see Supplementary Table 2 for detailed loadings of each task on each component). Component 1 accounted for 48.80% of the variance, with high loadings from WPV, WPPT, PPPT, OPN, NTD, and SN, and was therefore interpreted as the ‘General Semantics’ component. Component 2 (explaining 16.52% of the variance) loaded strongly on VP and SP, and was interpreted as the ‘Elementary Perception’ component. Component 3 (12.07% of the variance) showed a dominant loading on ALD, corresponding to the ‘Phonological Processing’ component.

**Figure 1.**
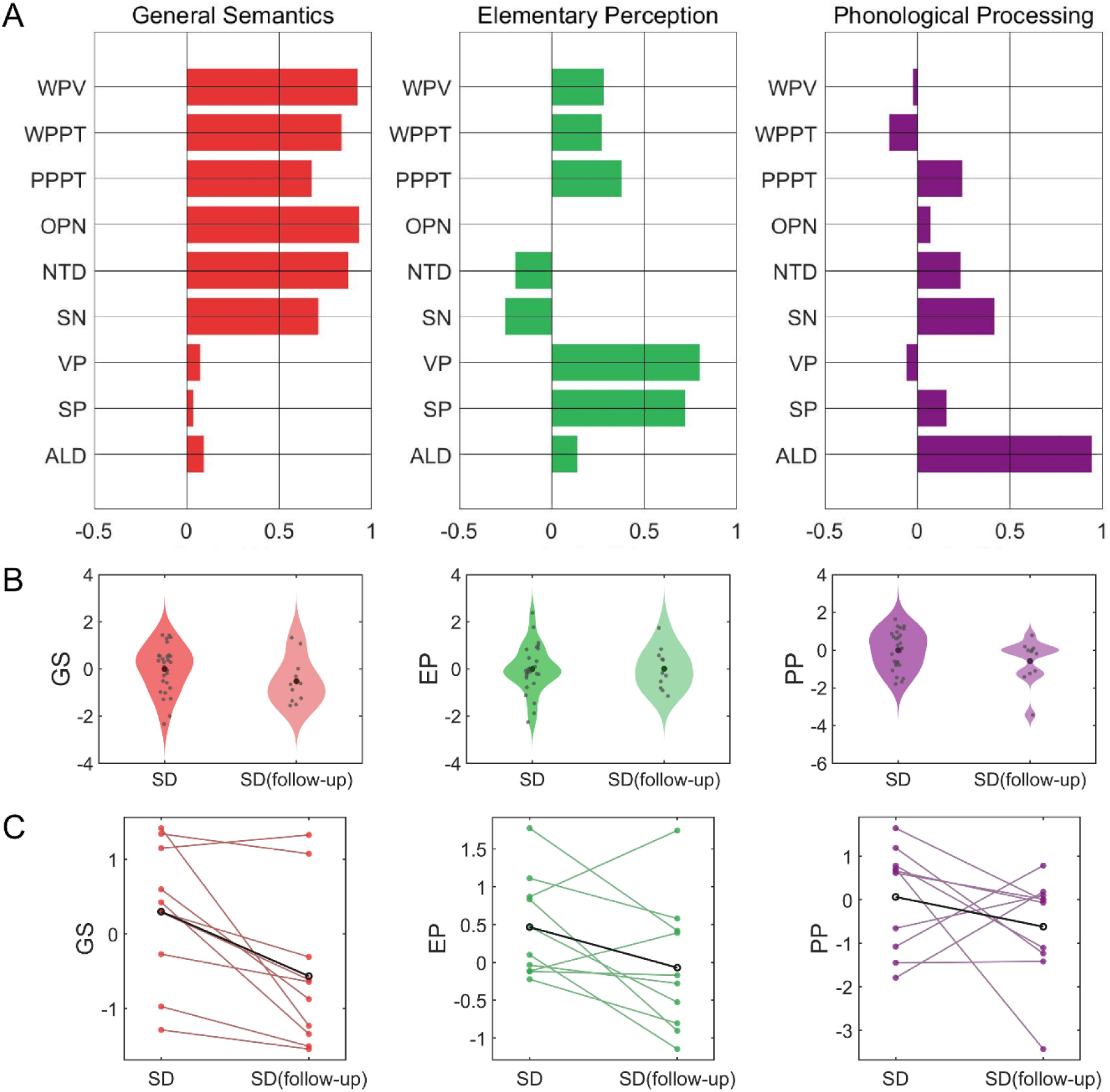
Principal component analysis and longitudinal profiles of three components in SDs. (A) The factor loadings from PCA. Three components with eigenvalues greater than 1 were identified: the first component, with high loadings on six general semantic tasks, was labeled ‘General Semantics’; the second, with high loadings on sound and visual perception tasks, was labeled ‘Elementary Perception’; and the third, mainly loaded on ALD, was labeled ‘Phonological Processing’. (B) Distribution of component scores at baseline and follow-up. (C) Longitudinal changes in individual patients’ component scores, shown as paired scatter plots with lines connecting baseline and follow-up data points. Abbreviations: WPV, Word-picture verification; WPPT, Word associative matching; PPPT, Picture associative matching; OPN, Oral picture naming; NTD, Naming to definition; SN, Oral sound naming; VP, Visual perception; SP, Sound perception; ALD, Auditory lexical decision.

Along with the component scores, we also analyzed changes over time in repetition performance, as shown in Figure 2. Overall, patients showed a declining trend across all assessed language domains—semantic processing, elementary perceptual abilities, phonological processing, and repetition—during the follow-up period, as illustrated in Figure 1B–C and 2.

**Figure 2.**
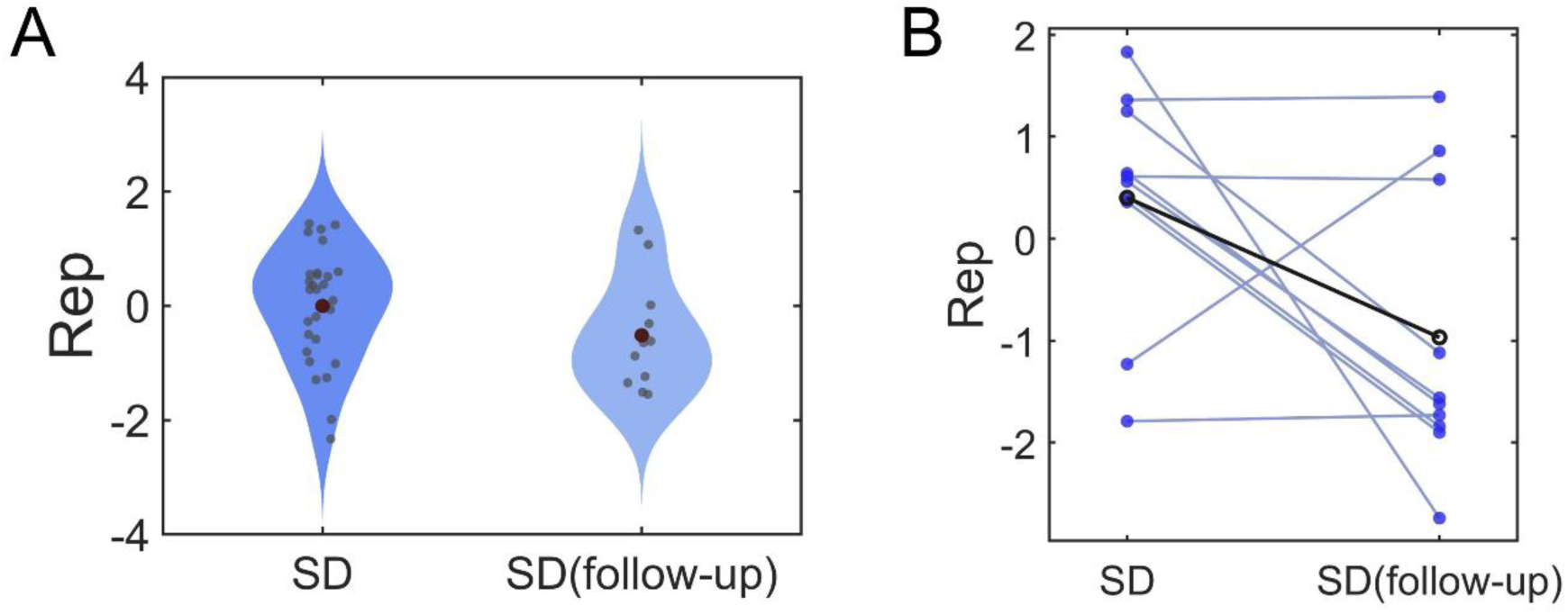
Longitudinal changes in repetition ability in SDs. (A) Violin plots showing the distribution of repetition task T-scores at baseline and follow-up. (B) Paired scatter plots with connecting lines illustrating within-subject changes in repetition scores across the two timepoints.

### GMV atrophy in SD patients

Figure 3 shows the atrophy of grey matter volume between HCs and SDs at baseline and follow-up (p < 0.01, FDR-corrected). Patients had the most severe atrophy in bilateral temporal lobes, insula, and ventral frontal lobes at baseline. Then, the atrophy in patients expanded to posterior temporal regions and frontal, parietal, and occipital areas at follow-up.

**Figure 3.**
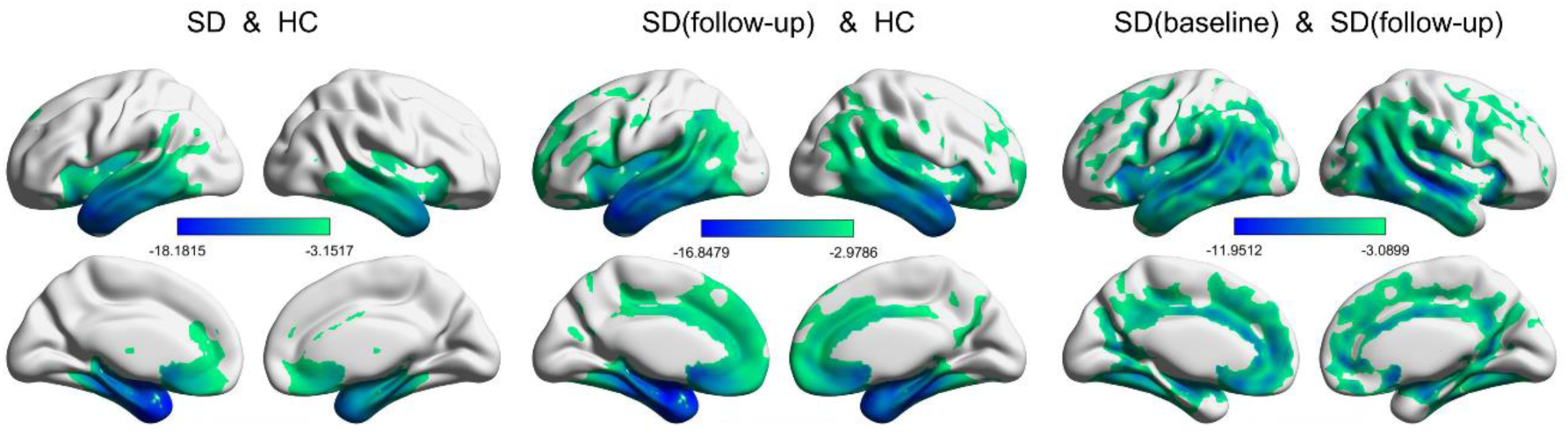
Brain atrophy map of semantic dementia. The figure highlights areas with significant differences in gray matter volume between HCs and SDs at baseline and follow-up (*p* < .01, FDR-corrected).

### The domain-segregation cortical language network dynamics in HCs

For each participant, frame-wise DCC produced 190 functional connectivity matrices (one per volume). By applying k-means clustering and the elbow criterion to all participants’ matrices (Nparticipants ∗ 190 frames), four reoccurring temporal states with distinct connectivity patterns and hub distributions were identified in HCs (Figure 4) and SDs at baseline and follow-up (Figure 4), which aligned with the “dynamic meta-networking framework of language.”

**Figure 4.**
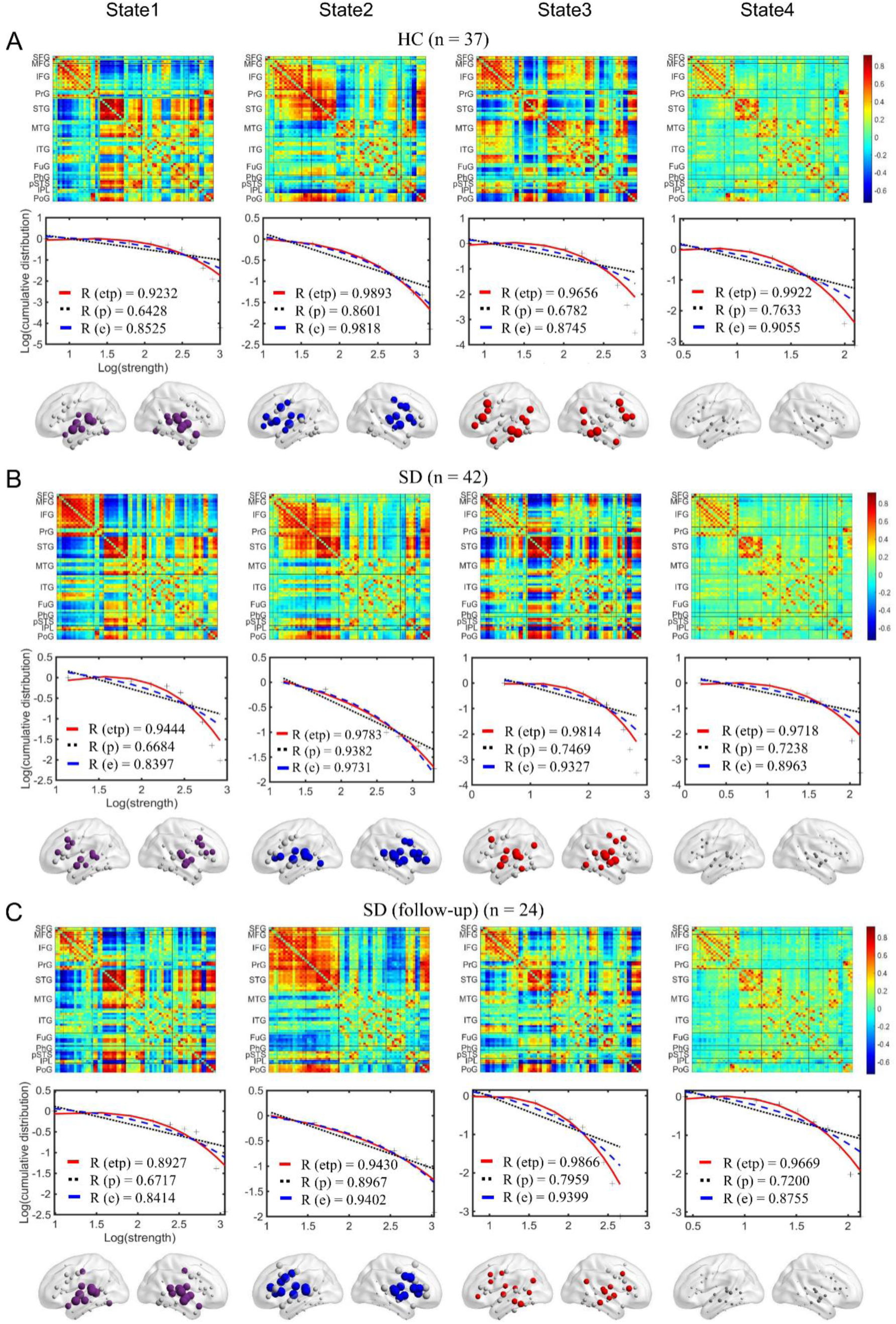
The four dFC states—State 1 (the central phonological state), State 2 (the dorsal articulatory state), State 3 (the ventral semantic state), and State 4 (the baseline state)—along with the corresponding cumulative nodal strength distributions and the state-dependent hub distributions in HCs (A) and SDs at baseline (B) and follow-up (C). In the log-log plots of the cumulative nodal strength distributions, the plus sign (black) represents observed data, the solid line (red) is the fit of the exponentially truncated power-law, *P*(*x*)∼*x*^*a*−1^exp (*x*/*x*^*c*^), the dashed line (blue) is an exponential, *P*(*x*)∼exp (*x*/*x*_*c*_), and the dotted line (black) is a power-law, *P*(*x*)∼*x*^*a*−1^. R² was calculated to assess the goodness-of-fit. In dFC states, warm color denotes positive connections, while cool color denotes negative connections. In the first three states, the first 20 nodes with the highest nodal strength were colored and defined as hubs.

In HCs, the central phonological state (State 1) was characterized by moderate-to-high positive connectivity between nodes in the bilateral STG, pSTS, and PoG, but by weak or moderate negative connectivity between prefrontal and temporal nodes. In the dorsal articulatory state (State 2), strong positive connectivity among prefrontal nodes and STG was observed. The ventral semantic state (State 3) distinguished itself from the central phonological and dorsal articulatory states by strong connectivity among inferior temporal nodes and the inferior parietal lobule (IPL). The baseline state (State 4) showed an overall weak connectivity pattern. The nodal strength distributions of all states were best fitted by an exponentially truncated power-law form, indicating the existence of a small set of highly connected hub nodes. The top 20 nodes with the highest nodal strengths were identified as hubs in each state (Figure 4). In the central phonological state, hubs were mainly located in the STG. In the dorsal articulatory state, hubs were primarily found in the prefrontal and superior temporal cortices. In the ventral semantic state, hubs were mostly present in the temporal cortex, posterior parietal cortex, and IFG. We did not consider hubs in the baseline state due to the overall weak connectivity. By calculating the spatial similarity between term-based activation and hub distribution, we found that hubs across the four states highly overlapped with activity related to speech perception, inner speech, phonological processing, and speech production, and partially overlapped with semantic processing (Figure 5). Given the critical role of STG in speech perception and phonological processing, we suspect hubs in the central phonological state are mainly involved in speech perception and phonological processing. Considering the importance of IFG and the ventral part of PrG in speech production, we hypothesize hubs in the dorsal articulatory state are primarily involved in speech production. Due to the key role of IPL, ATL, and the orbitofrontal cortex in semantic processing, we propose hubs in the ventral semantic state mainly support semantic processing. The baseline state may serve as a transition among these cognition-specific states.

**Figure 5.**
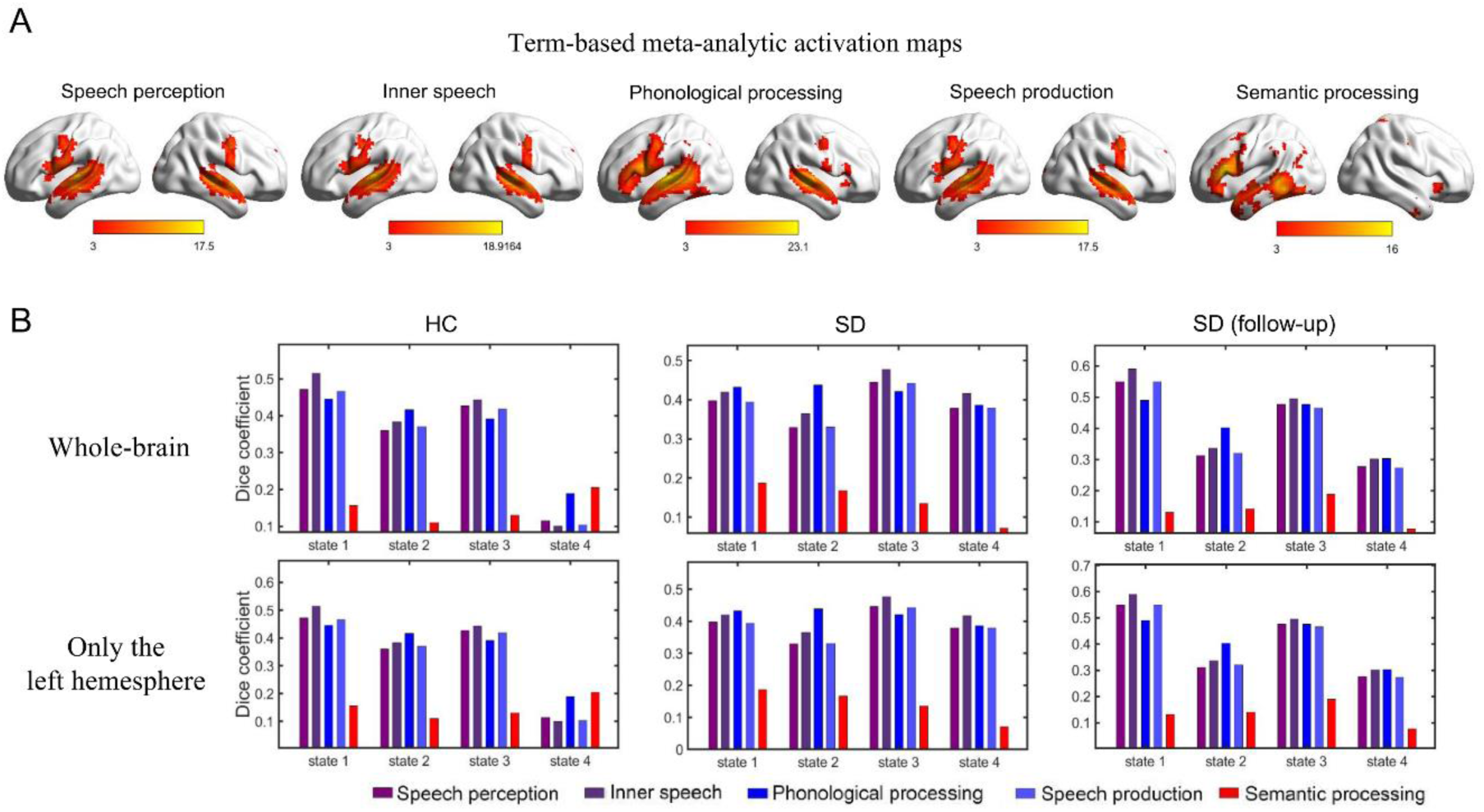
Functional relevance of hub distributions in language states. (A) The meta-analysis results for speech perception, inner speech, phonological processing, speech production, and semantic processing obtained from ‘NeuroQuery’ (https://neuroquery.org/). Each map was thresholded at *Z* = 3 (a typical threshold used by NeuroQuery) for illustration purposes, and only positive results are shown. (B) The Dice coefficients between binary images of rich-club nodes and meta results are presented. Due to the left-lateralized activations in the meta results, the Dice coefficients were calculated both across the whole brain and within the left hemisphere.

### Widespread and state-dependent disruptions in the cortical language network at baseline and follow-up

The cortical language network dynamics in SDs also showed four recurring temporal states (Figure 4). These four states exhibited high spatial similarity among HCs and SDs at both baseline and follow-up (Supplementary Figure 2). At baseline, although the cortical language network dynamics were severely disrupted in SDs, the nodal strength distributions across the four states were best described by an exponentially truncated power-law model. We defined the top 20 nodes with the highest nodal strength as hubs. In the central phonological state, hubs were located in the STG and the middle frontal gyrus (MFG). In the dorsal articulatory state, connections in MFG were weaker, but more hubs appeared in the posterior superior temporal gyrus (pSTG). The hub distribution in the ventral semantic state differed from HCs. Specifically, connections in the IFG, ATL, and ITG were significantly weakened, while hubs appeared in the STG and IPL. At follow-up, no significant differences were observed in connectivity patterns or hub distributions in the central phonological, dorsal articulatory, and baseline states. However, connections among the temporal cortex, posterior parietal cortex, and IFG all weakened in the ventral semantic state.

Statistical analyses indicated severe disruptions in the functional connectivity of all four states in SDs at both baseline and follow-up. Both hypo- and hyper-connectivity were observed (Figure 6), primarily driven by interhemispheric connections. Partial correlation analyses revealed that hypo-connectivity was positively associated with behavioral performance, whereas hyper-connectivity showed negative associations, suggesting dual effects of connectional diaschisis (Figure 6).

**Figure 6.**
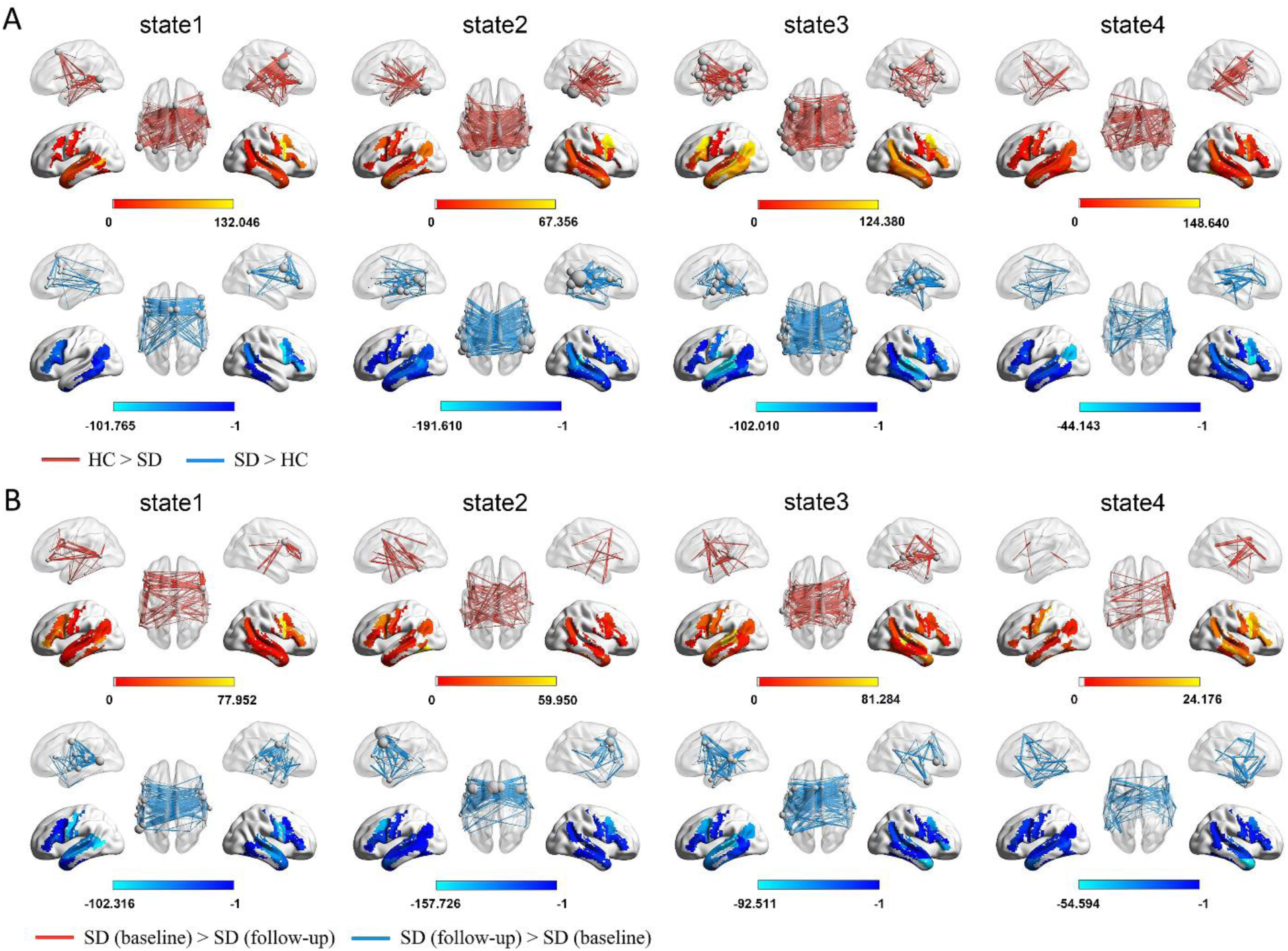
State-dependent hypo- and hyper-connectivity patterns in SDs at baseline (A) and follow-up (B) across the four dFC states: State 1 (the central phonological state), State 2 (the dorsal articulatory state), State 3 (the ventral semantic state), and State 4 (the baseline state). Edge *p*<0.01, component *p*<0.01 with NBS correction.

**Figure 7.**
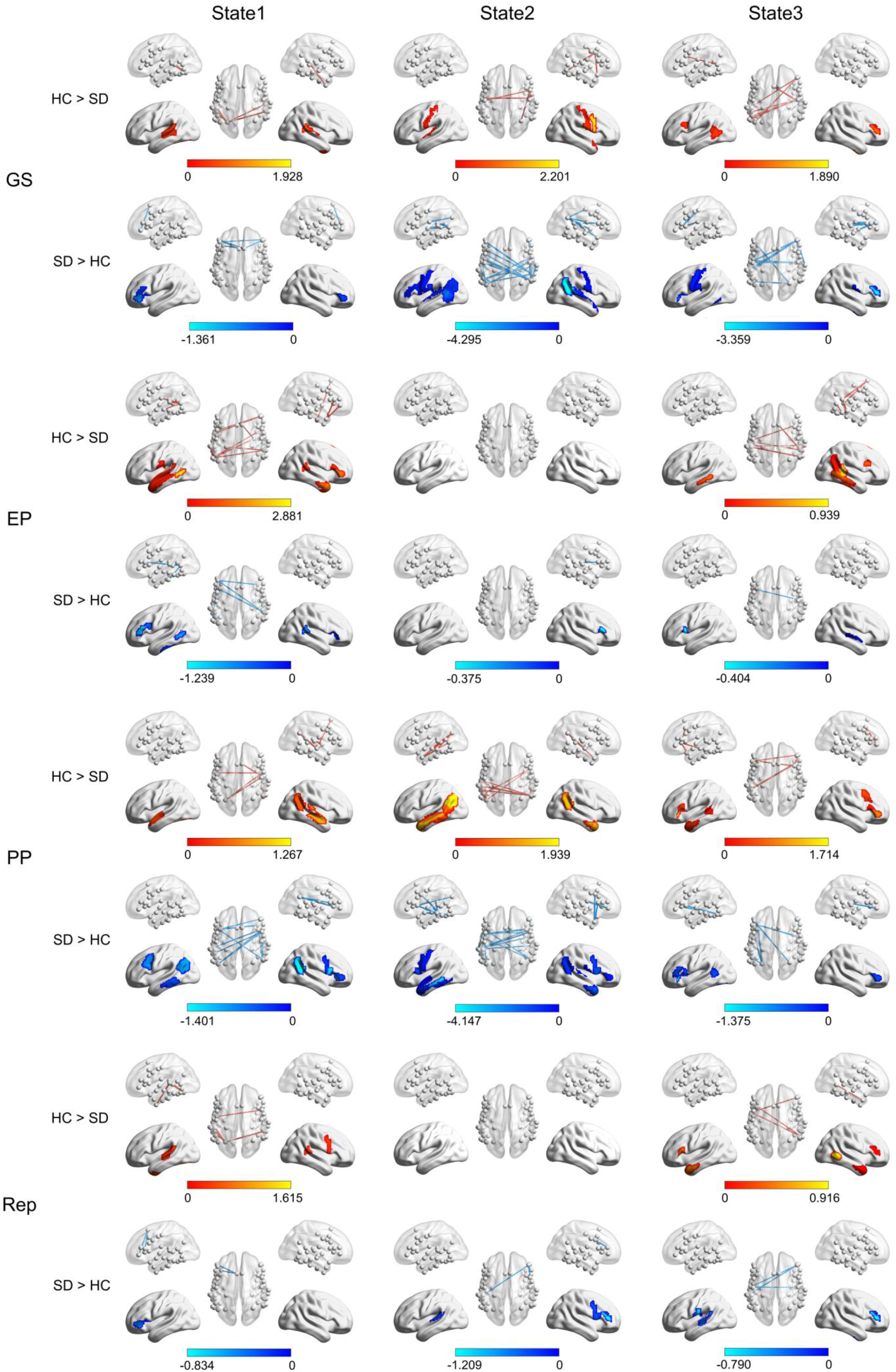
The partial correlation between network disruptions and four behavioral scores in SDs at baseline across three dFC states: State 1 (the central phonological state), State 2 (the dorsal articulatory state), and State 3 (the ventral semantic state). Edges indicate connections with significant correlation (*p* < 0.05, two-tailed, 10,000 permutations). Positive correlations are shown with red edges, representing increased connectivity linked to higher behavioral scores, while negative correlations are shown with blue edges, representing decreased connectivity linked to higher behavioral scores. Abbreviations: GS, General Semantics; EP, Elementary Perception; PP, Phonological Processing; Rep, Repetition.

Figure 8 illustrates the significant changes in nodal properties in SDs compared to HCs, with minimal differences observed between SDs at baseline and follow-up. In the central phonological state, there were notable reductions in nodal strength and nodal gE in the bilateral STG. In the dorsal articulatory state, a slight reduction of nodal strength was observed in the posterior ITG. In the ventral semantic state, there are significant increases in nodal strength, nodal gE, and nodal lE in the bilateral STG, along with significant decreases in nodal strength and nodal gE in the bilateral ITG and left IFG. However, partial correlation analyses revealed no significant associations between these nodal properties and behavioral performance, suggesting that nodal-level alterations may not directly map onto task outcomes.

**Figure 8.**
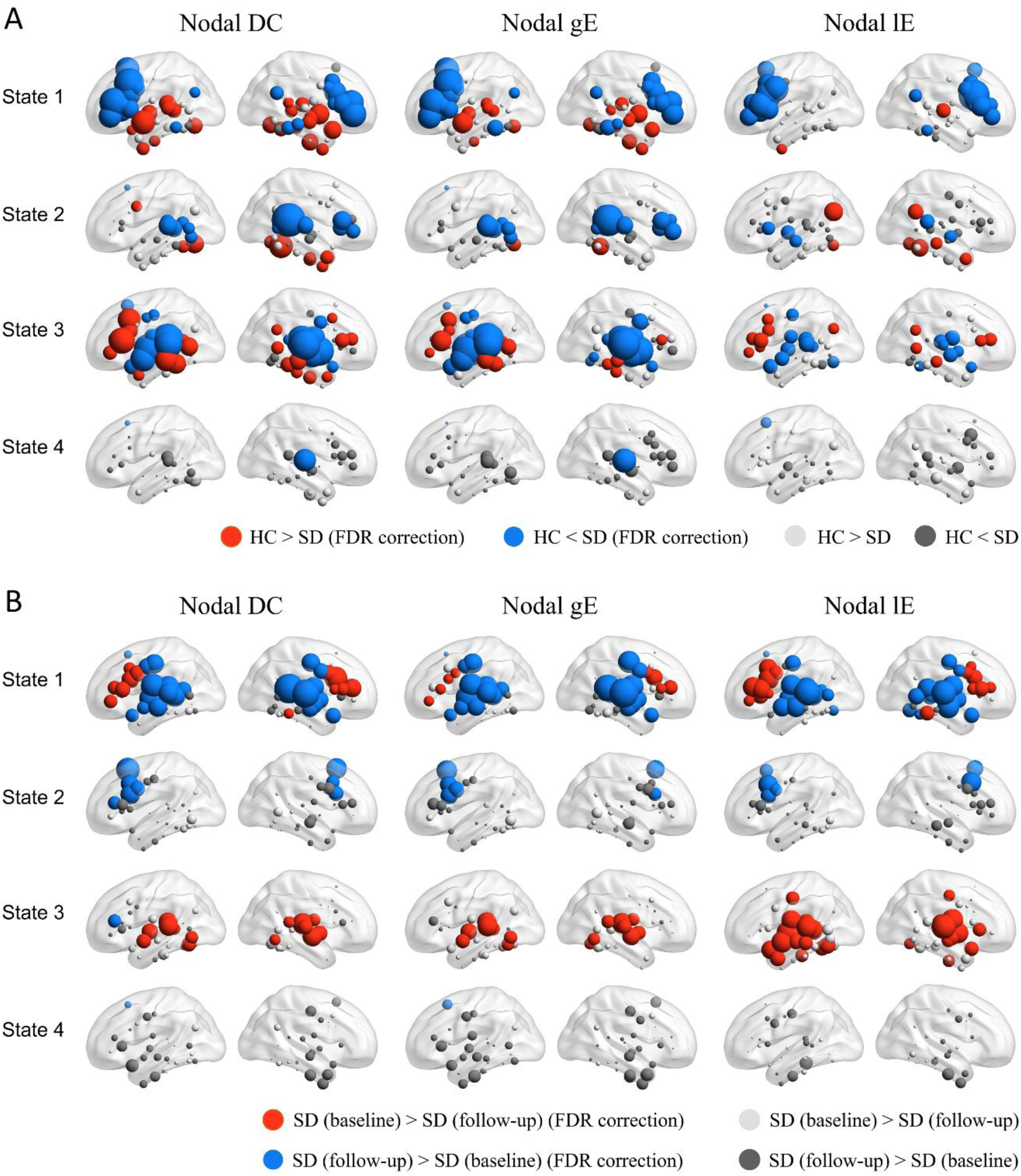
Changes in nodal properties in SD at baseline (A) and follow-up (B) across the four dFC states: State 1 (the central phonological state), State 2 (the dorsal articulatory state), State 3 (the ventral semantic state), and State 4 (the baseline state). Three nodal properties—nodal strength, nodal global efficiency (gE), and nodal local efficiency (lE)—were calculated for each participant by taking the median of state dFCs. Between-group statistical comparisons were conducted using two-sample t-tests (FDR correction, *p* < 0.05).

Figure 9 illustrates the changes in global properties in SDs compared to HCs and between SDs at baseline and follow-up. At baseline, the total functional connectivity strength of the central phonological state (uncorrected *p* < 0.001) and the ventral semantic state (uncorrected *p* < 0.01) significantly decreased compared to HCs. At follow-up, the gE and lE of the ventral semantic state both showed significant decreases (uncorrected *p* < 0.001). Partial correlation analyses were also performed between the three global topological properties and the four behavioral scores. The results indicated that none of the global properties had significant correlations with any of the behavioral domains, indicating no direct correspondence between global network changes and behavioral performance in SDs.

**Figure 9.**
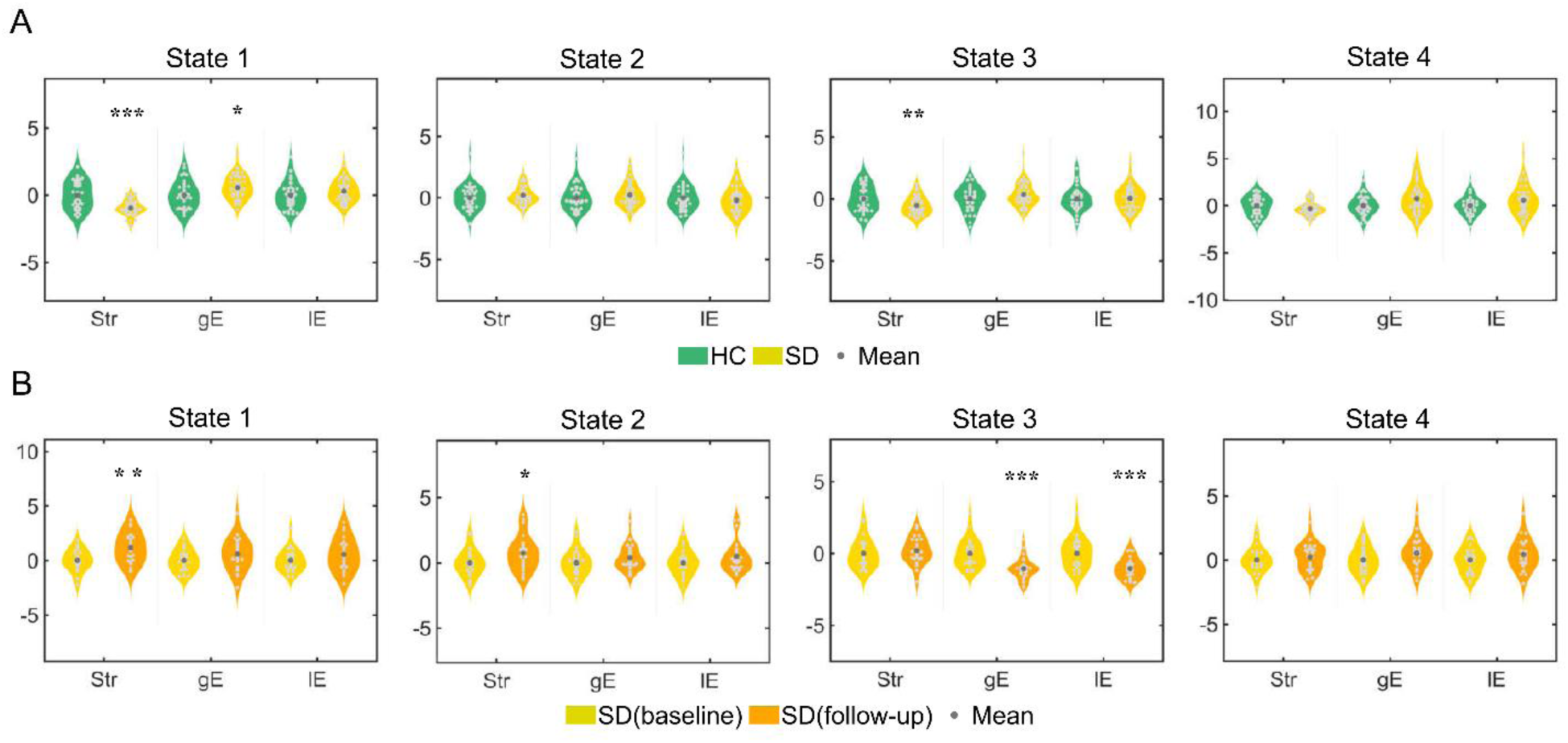
Changes in global properties of SD at baseline (A) and follow-up (B) across the four dFC states: State 1 (the central phonological state), State 2 (the dorsal articulatory state), State 3 (the ventral semantic state), and State 4 (the baseline state). Three network topological properties—total functional connectivity strength (Str), network global efficiency (gE), and local efficiencies (lE)—were calculated for each participant’s median dFC during each state. For illustration, the topological values were z-scored. Group differences in topological properties were assessed using multiple linear regression adjusted for covariates, followed by two-tailed t-tests to determine the significance of the regression coefficients (∗, *p* < 0.05; ∗∗, *p* < 0.01; ∗∗∗, *p* < 0.001).

### The dFCs significantly predicted each patient’s language scores

RVR-based dFC-linguistics prediction analyses revealed that dFCs of the central phonological state, dorsal articulatory state, ventral semantic state, baseline state, and the combinations of all four states predicted SDs’ GS (predicted vs. actual, *rs*: 0.07–0.47), EP (no significant correlation), and PP (*rs*: 0.37–0.60) (Figure 10). The model incorporating features from all four states showed the highest prediction accuracy. The dorsal articulatory state demonstrated the least predictive power. The score for ‘General Semantics’ was significantly predicted by the model including features of the central phonological state, ventral semantic state, baseline state, and the combined features of all four states. The ‘Phonological Processing’ score was strongly predicted by models using features from each individual state and the combined features, indicating robust predictive ability across both isolated and combined states.

**Figure 10.**
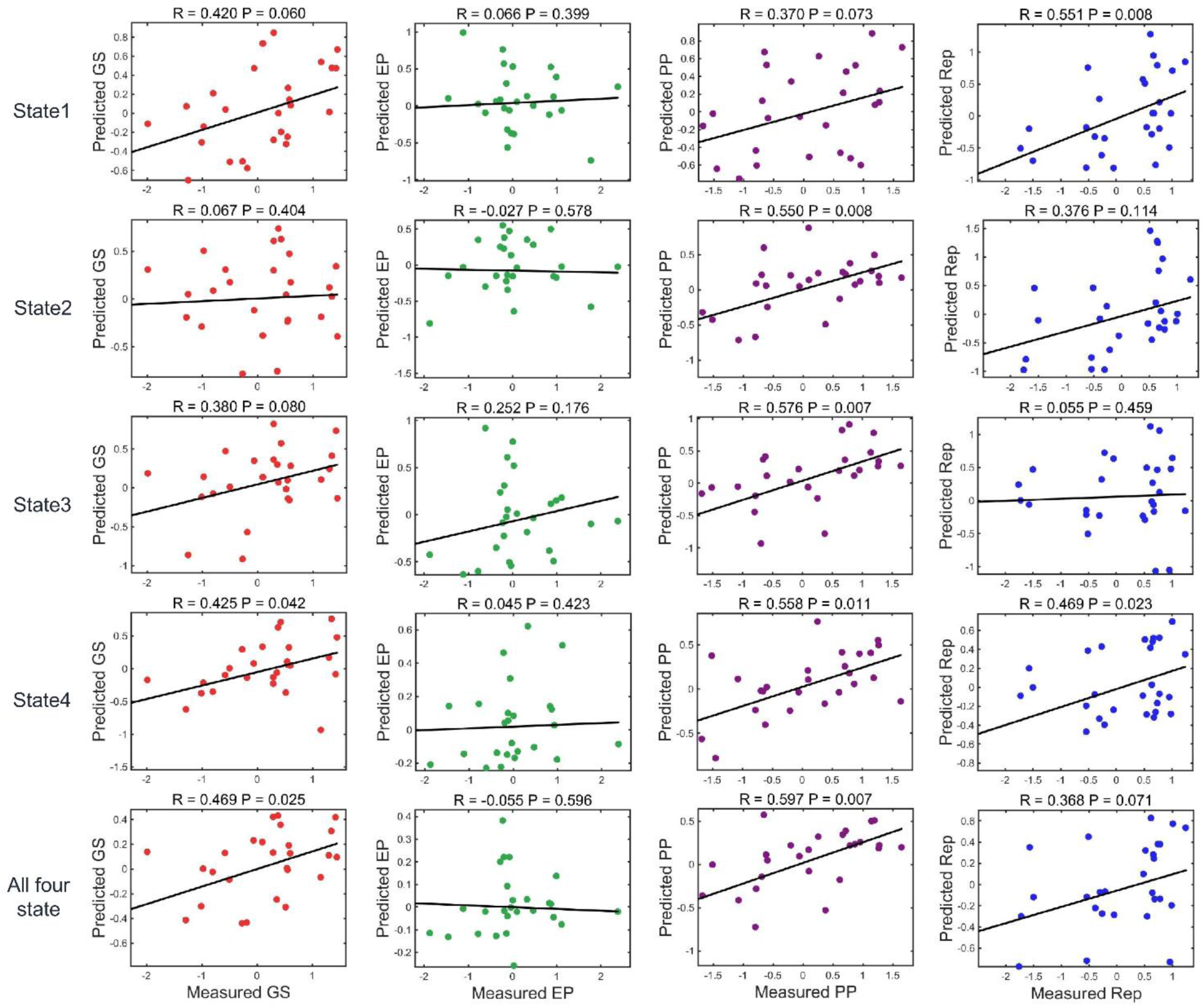
The accuracy and significance of the dFC-behavior model. The dFCs of the four states (State 1, the central phonological state; State 2, the dorsal articulatory state; State 3, the ventral semantic state; State 4, the baseline state) significantly predicted individual patients’ language deficits. The scatter plots displayed real (normalized within each group) and predicted scores for three factors, along with the corresponding linear fitted lines. The R-value represents the Pearson correlation coefficient between the predicted and actual values, and the model significance and *P*-values were derived from 1,000 permutation tests. Abbreviations: GS, General Semantics; EP, Elementary Perception; PP, Phonological Processing; Rep, Repetition.

## Discussion

Semantic dementia has long been considered a valuable pathological model for studying the neural basis of semantic representation; however, non-semantic linguistic impairments—such as phonological processing and speech repetition deficits—are often overlooked. In this study, we used a dynamic meta-networking framework of the cortical language network (Yuan, Xie, Wang, et al., 2023) to comprehensively map the network bases of cross-domain linguistic deficits. Behavioral assessments showed that SDs had significant impairments at baseline in general semantic ability, phonological processing, and speech repetition, all of which worsened over a two-year follow-up. In HCs, k-means clustering of dynamic functional connectivity patterns identified four recurring states of the language network, each featuring distinct hub distributions that reflected a domain-separation organizational pattern. Conversely, SDs showed disrupted domain-separation dynamics due to progressive brain atrophy, with both hypo- and hyperconnectivity across the four states. Notably, the ventral semantic state experienced the most severe disruption. Univariate analyses revealed that the state-specific hypo- and hyperconnectivity patterns were significantly linked to deficits across the three linguistic areas. Furthermore, machine learning-based predictions demonstrated that impairments in each domain could be specifically predicted by different states: semantic deficits were best predicted by the central phonological state, the ventral semantic state, and the baseline state; repetition deficits by the central phonological, the dorsal articulatory, and baseline states; and phonological deficits by all four states. Along with these connectivity disruptions, we observed state-specific changes in both nodal and global topological properties of the four meta states. However, these abnormal topological features did not significantly correlate with linguistic performance. Overall, these findings suggest that in SD, progressive atrophy spreads beyond the ATL to involve posterior temporal and frontal regions, leading to widespread dysfunction of the cortical language network and resulting in multi-domain linguistic impairments.

At baseline, SDs exhibited significant impairments across six semantic tasks (p < 0.001), with further decline observed at follow-up (p < 0.001). These behavioral deficits have long been associated with local atrophy and functional disconnection within the ventral language network. Previous structural MRI studies in SD consistently show pronounced atrophy in the bilateral anterior temporal lobes (Binney et al., 2010; Mummery et al., 2000), with additional degeneration in the insula, orbitofrontal cortex, posterior temporal, and lateral parietal regions as the disease progresses(Acosta-Cabronero et al., 2011; Agosta et al., 2010; Bejanin et al., 2020; Cousins et al., 2018). Our GMV analysis confirmed these findings, revealing significant grey matter loss in the bilateral ATL at baseline, which extended to posterior temporal, parietal, and occipital regions at follow-up. Functional connectivity research using static functional connectivity (sFC) and task-based fMRI has consistently shown widespread disruption of the semantic network in SD. Previous studies reported intra- and inter-hemispheric connectivity within the ATL, ITG, and angular gyrus (AG) —key hubs of the semantic system (Chen et al., 2022; Guo et al., 2013). However, static approaches cannot capture the transient and flexible nature of network reconfigurations within language network and tend to underestimate the flexible segregation and integration of ventral and dorsal streams involved in language and speech processing (Fridriksson et al., 2016; Hickok, 2022; Rauschecker & Scott, 2009; Ries et al., 2019). The dynamic meta-networking framework we adopted offers a more comprehensive approach to map domain-specific network disruptions and cross-domain network alterations resulting from progressive brain atrophy.

Our dynamic meta-networking analyses identified four temporally-resolved functional states, each characterized by distinct domain-related hub distributions. These domain-separation cortical language dynamics were disrupted by the progressive brain atrophy of SD. Given the pronounced atrophy in the ATL, the most immediate disruption was observed in the ventral semantic state, where ATL served as a core hub supporting semantic integration. In addition, our VBM results revealed posterior temporal atrophy at both baseline and follow-up, with greater progression at follow-up, leading to substantial disruption in the central phonological state, which is anchored in the superior temporal gyrus. Both the central phonological state and the ventral semantic state are centered on temporal lobe regions, reflecting the breakdown of the ventral semantic unification pathway and the central phonological stream. Although the prefrontal cortex did not exhibit pronounced structural atrophy at either timepoint, the dorsal articulatory state—associated with the dorsal articulatory pathway—also showed substantial disruption. This likely reflects the functional interdependence between the dorsal stream and the other two pathways (Fedorenko & Thompson-Schill, 2014; Hickok & Poeppel, 2007; Saur et al., 2008). These results indicate that while the three pathways showed domain-specific segregation during resting-state dynamics, functional interactions persisted, as evidenced by the partially shared hub distributions across states (Yuan, Xie, Wang, et al., 2023).

Semantic impairment is the key linguistic deficit in SDs, marked by significant deficits at baseline and further decline over time. In the ventral semantic state, core hubs in the ATL and ITG showed substantial atrophy. Topological analyses also revealed reductions in both global and local efficiency, along with decreased nodal centrality in bilateral ITG and the IFG. These deficits indicate a breakdown of the ventral semantic pathway, which is widely recognized as crucial for semantic representation and integration (Binney et al., 2010; Huang et al., 2023; Ralph et al., 2017). This pathway has been confirmed repeatedly using multimodal imaging and lesion network mapping techniques (Chen et al., 2017; Y. Zhao et al., 2017). Machine learning models showed that semantic impairment was predicted not only by the ventral semantic state but also by the central phonological state and baseline state. This suggests functional interactions across states and indicates that semantic deterioration in SD involves not only disruptions in the ventral semantic pathway but also its interaction with phonological perception processes and baseline cortical activity. In contrast, the dorsal articulatory state did not predict semantic deficits, reinforcing the functional separation between ventral semantic and dorsal articulatory pathways (Hickok, 2022; Hickok & Poeppel, 2007).

Phonological processing was also impaired at baseline and showed a gradual worsening, although at a less steep rate than the decline in semantic processing. Consistent with previous research, speech perception and phonological processing are known to depend on a distributed network centered around the STG, AG, and Wernicke’s area (Jefferies et al., 2006, 2011). Recent neurocomputational models have further clarified the role of the STG as a key site for extracting multiple speech features, including phonemic identity, prosody, and temporal structure (Bhaya-Grossman & Chang, 2022). Our VBM analysis confirmed these findings, showing significant atrophy in these regions. Likewise, dynamic network analysis revealed notable changes in the topological properties of the central phonological state. Machine learning models indicated that phonological performance was predicted not only by the central phonological state but also by the dorsal articulatory state and the ventral semantic state. This could be due to two reasons. First, although the auditory lexical decision task mainly aims to test phonological processing, it also involves articulatory rehearsal and semantic plausibility assessments. Evidence for articulatory involvement is seen in overlapping activation in the left pSTG during both speech perception and production tasks (Buchsbaum et al., 2001). On the semantic front, evidence suggests that even implicit semantic judgments can help facilitate lexical decisions, as shown by overlapping phonological and semantic activations in temporal regions (Vigneau et al., 2006). Second, the hub distributions of the central phonological state, the dorsal articulatory state, and the ventral semantic state partially overlap—especially within the posterior temporal areas—implying that interactions among these states may work together to support phonological processing.

Speech repetition performance in SDs showed a significant yet relatively mild impairment at baseline (*p* = 0.033) and a progressive decline over time. Previous studies indicated that repetition abilities, especially for simple words and phrases, are often intact in the early to moderate stages of SD (Gorno-Tempini et al., 2011; Mesulam et al., 2012). This subtle but detectable deterioration aligns with findings from Jefferies et al. (2006), which showed that both semantic and phonological impairments can independently contribute to deficits in delayed auditory repetition, depending on the location of cortical degeneration. Importantly, resting-state network analysis suggested that the dorsal articulatory state—linked to articulatory planning and motor execution—remained relatively stable compared to other states. Speech repetition performance was significantly predicted by the central phonological state, the dorsal articulatory state, and the baseline state, but not by the ventral semantic state, indicating that repetition mainly relies on the dorsal and central processing streams rather than semantic pathways.

Elementary perceptual functions, assessed through simple visual and sound perception tasks, showed preserved visual performance but mild deficits in sound perception in SDs. Importantly, none of the four states significantly predicted perceptual performance, highlighting the domain specificity of the dynamic meta-networking model. Mild auditory impairments (*p* < 0.05) may indicate involvement of secondary auditory regions, such as the posterior STG.

## Conclusion

By applying a domain-separation meta-networking framework to the cortical language network, we identified the network-level foundations of multi-domain linguistic decline in semantic dementia. Progressive atrophy starting in the anterior temporal lobe directly disrupted the ventral semantic processing stream. As the atrophy extended to the posterior temporal cortex and prefrontal regions, both the central stream responsible for phonological processing and the dorsal stream involved in speech production were increasingly affected. As a result, the domain-segregation and integration functions of the entire cortical language network were significantly impaired. These findings emphasize the crucial role of the language system’s domain-segregated yet interconnected architecture and highlight the importance of early intervention in SDs to potentially slow the overall deterioration of their language network.

## Supporting information

Supplementary Methods. Behavioral tasks. Supplementary Methods. Relevance Vector Regression (RVR) algorithm. Supplementary Figure 1. Language networ

## Conflict of Interest

No competing financial interests exist.

## Data and Code Availability Statement

All data and study materials are available from the corresponding author on reasonable request. The DCC toolbox was available at https://github.com/yuanbinke/Naturalistic-Dynamic-Network-Toolbox.

## Funding

The study is supported by the National Social Science Foundation of China (20&ZD296), Key-Area Research and Development Program of Guangdong Province (2019B030335001), National Natural Science Foundation of China (32400862), Research Center for Brain Cognition and Human Development, Guangdong, China (2024B0303390003), Shanghai Medical Innovation and Development Foundation “Brain Health Youth Fund - Precision Diagnosis and Treatment Research on Alzheimer’s Disease”(SMIDF-150-2025A30), Basic Scientific Research Project of Shanghai Sixth People’s Hospital (ynqn202222), Special Project for Clinical Research of Shanghai Municipal Health Commission (202440009), National Natural Science Foundation of China (82501892).

## Notes

### Competing Interest Statement

The authors have declared no competing interest.

