## Supplementary Methods. Behavioral tasks. Supplementary Methods. Relevance Vector Regression (RVR) algorithm. Supplementary Figure 1. Language networ for "Beyond the Anterior Temporal Lobe: Domain-Related Degeneration of Cortical Language Network Dynamics in Semantic Dementia"

**Supplementary Material**

**Supplementary Methods**

**Behavioral tasks**

We designed six general semantic tasks and six modality-specific semantic tasks as follows. The general semantic tasks examined the general aspects of semantic knowledge with various modalities of input and output.

***General semantic tasks***

Oral picture naming. Subjects were instructed to name each object whose picture was presented on the screen. The first complete response was scored for each item.

Oral sound naming. Subjects were instructed to speak out the name of the object upon hearing the target sound through earphones.

Picture associative matching. This task had the same format as the Pyramids and Palm Trees Test, with each trial containing three photographed objects on the screen. Subjects needed to judge which of the two bottom photographs was semantically closer to the top photograph.

Word associative matching. This task was the same as the Picture associative matching task, but the pictures were replaced with Chinese written names.

Word-picture verification. Participants were required to decide whether the name and the picture matched.

Naming to definition. Participants were asked to name the object whose definition was visually and aurally presented.

**Details of the Relevance Vector Regression (RVR) Algorithm**

RVR is a Bayesian framework for learning sparse regression models. In RVR, only some samples (smaller than the training sample size), termed the “relevance vectors,” are used to fit the model:

$$y\left( x \right)=\sum_{i=1}^{m} \omega_{i}\tau_{i}+\epsilon,$$

where $\tau_{i}$are basis functions, and $\epsilon$ is normally distributed with mean 0 and variance $\beta$. RVR uses training data to build a regression model:

$$y=\theta\omega+\epsilon,$$

where $y={[y_{1},y_{2},\ldots y_{n}]}^{T},\theta=\left[ \theta_{1},\theta_{2},\ldots\theta_{n} \right],\theta_{i}={[\tau_{i}\left( \chi_{1} \right),\tau_{i}\left( \chi_{2} \right),\ldots\tau_{i}\left( \chi_{n} \right)]}^{T}.$Each vector$\theta_{i}$, consisting of the values of the basis function $\tau_{i}$ for the input vectors, is a relevance vector.

The model parameters $\beta$ were found by using the maximum likelihood estimates from the conditional distribution: $p\left( y | \alpha,\beta\right)=\mathcal{N}\left( y | 0,C \right),$ where the $C=\beta I_{n}+\Phi A^{-1}\Phi^{T}.$ To make the RVR favor sparse regression models, prior distributions were assumed for both $\omega_{i}$ and $\beta^{-1}$, i.e. $p\left( \omega_{i} | \alpha_{i} \right)\mathcal{=N}(0,{\alpha_{i}}^{-1}$). The ways of treating priors, however, lead to the same relevance vector machine construction.

**Supplementary Figures**

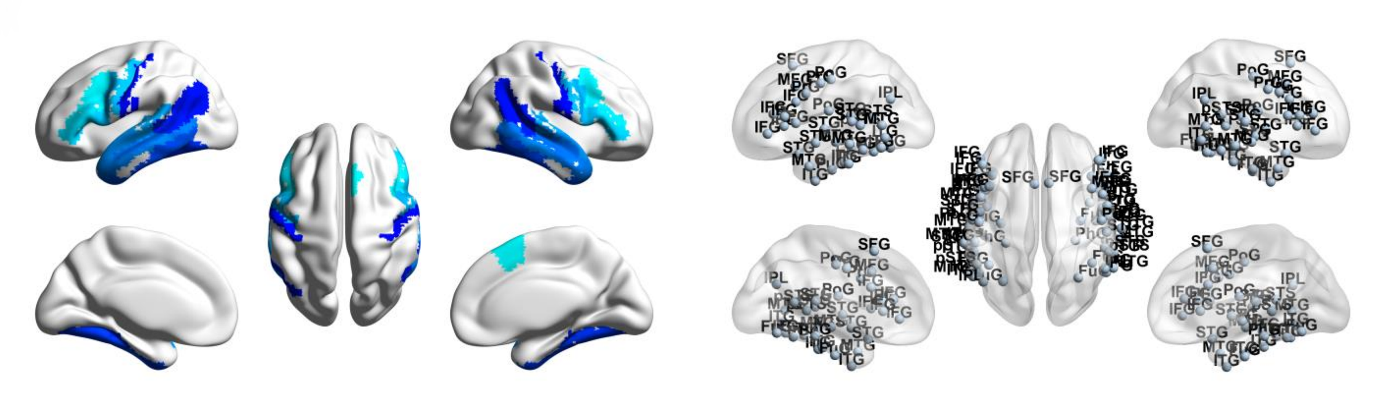

**Supplementary Figure 1.** The language network, consisting of 68 regions involved in language processing, was selected from the Brainnetome Atlas (n = 246, <https://atlas.brainnetome.org/bnatlas.html>) (Fan et al., 2016), according to each node’s meta-analytic behavioral domains and paradigm classes.

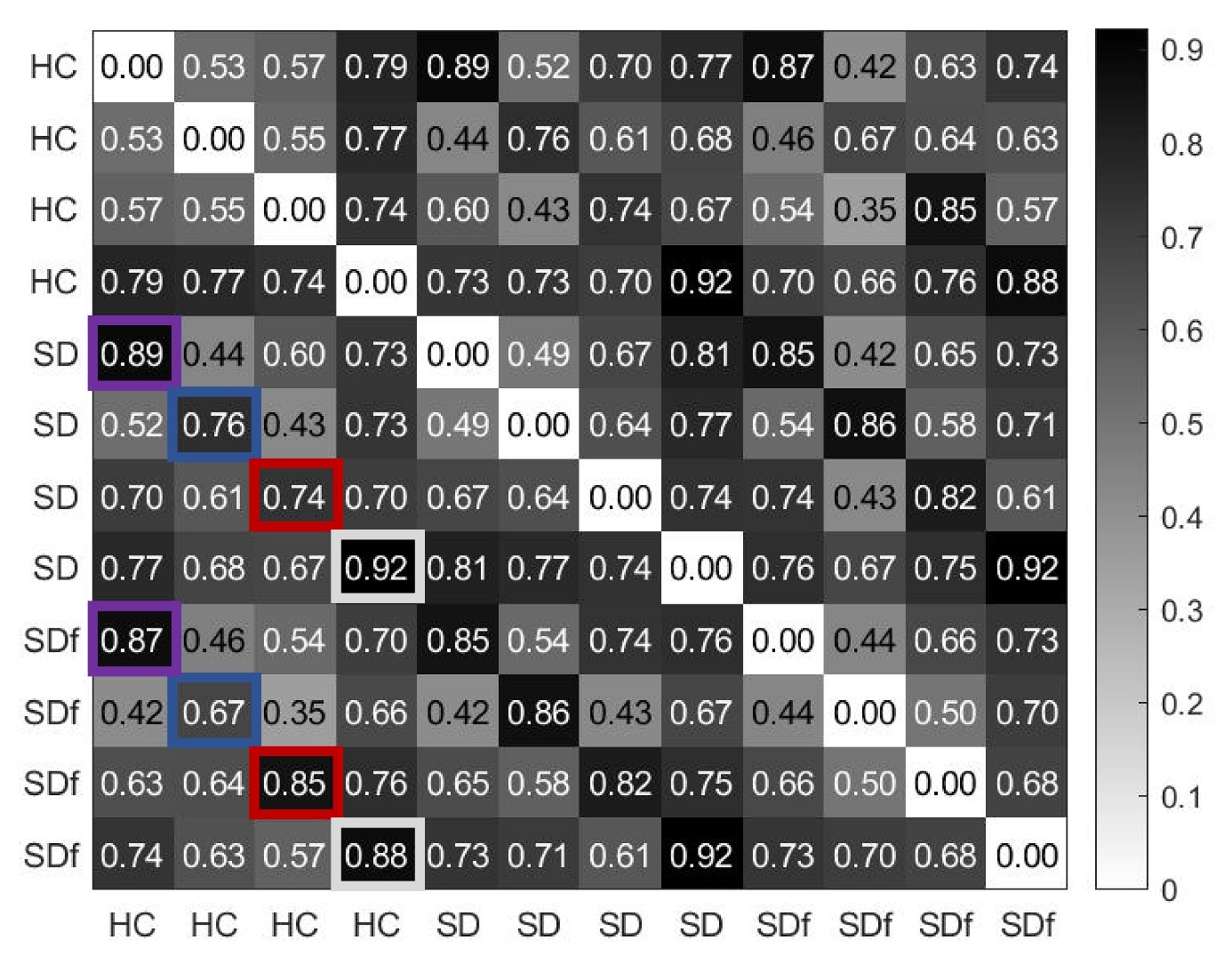

**Supplementary Figure 2.** The spatial correlation coefficients among the four states of HCs and

SDs at baseline (SD) and follow-up (SDf).

**Supplementary tables**

Supplementary Table 1. Anatomical regions，language-related behavioral domains, and paradigm classes of the language network. Coordinates are in the standard Montreal Neurologic Institute Space.

| ID | x | y | z | anatomy | behavioral domains | paradigm classes |
| --- | --- | --- | --- | --- | --- | --- |
| 1 | -4.6 | 15.7 | 53.2 | SFG | Execution. Speech, Phonology, Semantics, Speech | Word Generation (Covert and Overt) |
| 2 | 7.0 | 16.4 | 54.7 | SFG | The homolog of node 1 | - |
| 17 | -41.9 | 13.6 | 36.4 | MFG | Phonology, Semantics | Semantic. Monitor/Discrimination, Word Generation (Covert) |
| 18 | 42.2 | 11.9 | 38.4 | MFG | The homolog of node 17 | - |
| 29 | -45.6 | 13.0 | 23.4 | IFG | Phonology, Semantics, Speech, and Syntax | Phonological. Discrimination, Semantic. Monitor/Discrimination |
| 30 | 45.0 | 15.7 | 25.0 | IFG | The homolog of node 29 | - |
| 31 | -47.6 | 31.6 | 13.6 | IFG | Phonology, Semantics, Speech, and Syntax | Phonological. Discrimination, Semantic. Monitor/Discrimination, Word Generation (Covert and Overt) |
| 32 | 47.8 | 35.1 | 13.2 | IFG | The homolog of node 31 | - |
| 33 | -52.2 | 22.5 | 11.4 | IFG | Semantics, Speech, and Syntax | Reading (Covert), Semantic. Monitor/Discrimination, Word Generation (Covert and Overt) |
| 34 | 54.2 | 23.9 | 11.8 | IFG | The homolog of node 33 | - |
| 35 | -49.1 | 36.3 | -2.8 | IFG | Semantics, Speech, and Syntax | Semantic. Monitor/Discrimination, Word Generation (Covert) |
| 36 | 50.8 | 36.8 | -0.7 | IFG | The homolog of node 35 | - |
| 37 | -39.5 | 22.9 | 3.7 | IFG | Phonology, Semantics, Speech, and Syntax | Semantic. Monitor/Discrimination, Word Generation (Covert) |
| 38 | 42.1 | 22.0 | 3.2 | IFG | The homolog of node 37 | - |
| 39 | -51.3 | 13.2 | 6.0 | IFG | Phonology, Semantics, Speech | Music. Comprehension/Production, Recitation/Repetition. (Covert), Word Generation (Covert) |
| 40 | 53.6 | 14.3 | 11.8 | IFG | The homolog of node 39 | - |
| 53 | -49.5 | -7.1 | 38.8 | PrG | Execution. Speech | Reading (Overt), Recitation/Repetition. (Overt) |
| 54 | 54.9 | -2.0 | 33.3 | PrG | Execution. Speech | Reading (Overt), Recitation/Repetition. (Overt) |
| 63 | -49.1 | 4.7 | 30.5 | PrG | Orthography, Phonology, Semantics, Speech, and Syntax | Phonological. Discrimination, Reading (Covert) |
| 64 | 51.1 | 7.2 | 30.9 | PrG | The homolog of node 63 | - |
| 71 | -53.7 | -32.1 | 12.4 | STG | Execution. Speech, Phonology, and Speech | Music. Comprehension/Production, Passive Listening, Phonological. Discrimination, Reading (Overt), Recitation/Repetition. (Covert and Overt) |
| 72 | 54.5 | -23.7 | 10.6 | STG | Execution. Speech, Phonology | Music. Comprehension/Production, Passive Listening, Phonological. Discrimination, Recitation/Repetition. (Overt) |
| 73 | -50.1 | -10.3 | 1.1 | STG | Execution. Speech, Phonology, Speech | Music. Comprehension/Production, Passive Listening, Phonological. Discrimination, Reading (Overt), Recitation/Repetition. (Overt) |
| 74 | 51.1 | -3.7 | -0.9 | STG | Execution. Speech, | Music. Comprehension/Production, Passive Listening, Recitation/Repetition. (Overt) |
| 75 | -62.8 | -32.8 | 7.4 | STG | Execution. Speech, Phonology, Semantics, Speech | Passive Listening, Phonological. Discrimination, Reading (Overt), Semantic. Monitor/Discrimination |
| 76 | 66.5 | -20.8 | 6.6 | STG | Execution. Speech, Phonology, Speech | Music. Comprehension/Production, Passive Listening, Phonological. Discrimination, Reading (Overt) |
| 77 | -45.1 | 10.5 | -19.4 | STG | The homolog of node 78 | - |
| 78 | 47.1 | 12.3 | -19.7 | STG | Speech | Film Viewing, Passive Listening |
| 79 | -55.1 | -3.2 | -10.1 | STG | Execution. Speech, Phonology, Semantics, Speech | Music. Comprehension/Production, Passive Listening, Phonological. Discrimination, Reading (Overt) |
| 80 | 55.8 | -12.5 | -5.2 | STG | Execution. Speech, Phonology, Semantics, Speech | Music. Comprehension/Production, Passive Listening, Phonological. Discrimination, Reading (Overt), Semantic. Monitor/Discrimination |
| 81 | -65.2 | -30.9 | -11.3 | MTG | Semantics | Semantic. Monitor/Discrimination |
| 82 | 64.5 | -29.2 | -13.2 | MTG | The homolog of node 81 | - |
| 83 | -53.2 | 2.2 | -29.6 | MTG | Language | Semantic. Monitor/Discrimination, Reading (Covert) |
| 84 | 51.1 | 5.7 | -31.8 | MTG | Language | Passive Listening |
| 85 | -58.9 | -57.6 | 4.3 | MTG | Semantics and Syntax | Semantic. Monitor/Discrimination, Word Generation (Overt) |
| 86 | 60.1 | -53.3 | 2.9 | MTG | The homolog of node 85 | Film Viewing |
| 87 | -58.5 | -19.8 | -9.4 | MTG | Phonology, Semantics, Speech, and Syntax | Passive Listening, Phonological. Discrimination, Reading (Covert), Semantic. Monitor/Discrimination |
| 88 | 58.3 | -15.4 | -10.1 | MTG | Semantics and Speech | Passive Listening, Phonological. Discrimination |
| 89 | -45.5 | -26.7 | -26.1 | ITG | Orthography and Semantics | Reading (Covert), Semantic. Monitor/Discrimination |
| 90 | 45.8 | -14.6 | -32.4 | ITG | The homolog of node 89 | - |
| 91 | -50.5 | -57.0 | -14.1 | ITG | Phonology and Semantics | Naming (Overt) |
| 92 | 53.5 | -52.4 | -18.5 | ITG | Phonology and Semantics | - |
| 93 | -43.7 | -2.9 | -41.4 | ITG | Semantics | Semantic. Monitor/Discrimination |
| 94 | 40.5 | -2.9 | -41.4 | ITG | The homolog of node 93 | - |
| 97 | -55.2 | -60.3 | -6.0 | ITG | Semantics and Speech | Film Viewing, Naming (Overt) |
| 98 | 54.2 | -56.9 | -8.6 | ITG | The homolog of node 97 | - |
| 99 | -58.8 | -42.1 | -16.0 | ITG | Orthography and Semantics | Naming (Overt), Reading (Covert), Semantic. Monitor/Discrimination, Word Generation (Overt) |
| 100 | 60.5 | -40.5 | -17.1 | ITG | The homolog of node 99 | - |
| 101 | -54.9 | -30.3 | -27.4 | ITG | Semantics | - |
| 102 | 53.7 | -30.3 | -26.3 | ITG | The homolog of node 101 | - |
| 103 | -32.4 | -16.6 | -32.3 | FuG | Semantics and Speech | Naming (Overt), Semantic. Monitor/Discrimination |
| 104 | 33.1 | -14.6 | -34.1 | FuG | Semantics | Naming (Overt), Semantic. Monitor/Discrimination |
| 105 | -30.6 | -64.4 | -14.1 | FuG | Orthography, Semantics, Speech | Naming (Covert and Overt) |
| 106 | 31.3 | -61.4 | -13.7 | FuG | Language | Naming (Covert and Overt) |
| 107 | -42.3 | -50.9 | -17.3 | FuG | Orthography, Phonology, Semantics, Speech | Naming (Covert and Overt), Phonological. Discrimination, Reading (Covert), and Semantic. Monitor/Discrimination |
| 108 | 42.7 | -49.1 | -18.6 | FuG | Orthography, Semantics | Naming (Covert) |
| 113 | -28.3 | -32.6 | -16.9 | PhG | Semantics | Naming (Overt), Semantic. Monitor/Discrimination |
| 114 | 28.8 | -30.7 | -17.5 | PhG | The homolog of node 113 | Passive Listening, Semantic. Monitor/Discrimination |
| 121 | -54.4 | -39.8 | 4.2 | pSTS | Phonology, Semantics, Speech, Syntax | Passive Listening, Phonological. Discrimination, Reading (Covert), Semantic. Monitor/Discrimination, Word Generation (Covert and Overt) |
| 122 | 52.9 | -36.8 | 3.1 | pSTS | Execution. Speech, Phonology, Semantics, and Speech | Passive Listening, Phonological. Discrimination, Reading (Overt) |
| 123 | -52.4 | -50.3 | 10.8 | pSTS | Orthography, Semantics, Speech, and Syntax | Reading (Covert), Semantic. Monitor/Discrimination |
| 124 | 56.5 | -40.1 | 12.5 | pSTS | The homolog of node 123 | Passive Listening |
| 143 | -46.8 | -64.7 | 25.8 | IPL | Language | Semantic. Monitor/Discrimination, |
| 144 | 53.0 | -54.1 | 24.4 | IPL | The homolog of node 143 | - |
| 155 | -50.4 | -15.8 | 42.1 | PoG | Execution. Speech | Recitation/Repetition. (Overt) |
| 156 | 50.3 | -14.2 | 43.7 | PoG | Execution. Speech | Reading (Overt), Recitation/Repetition. (Overt) |
| 157 | -55.8 | -14.0 | 16.2 | PoG | Execution. Speech | Recitation/Repetition. (Overt) |
| 158 | 55.2 | -10.2 | 15.0 | PoG | Execution. Speech | Recitation/Repetition. (Overt) |

*SFG*, superior frontal gyrus; *MFG*, middle frontal gyrus; *IFG*, inferior frontal gyrus; *OrG*, orbital gyrus; *PrG*, precentral gyrus; *STG*, superior temporal gyrus; *MTG*, middle temporal gyrus; *ITG*, inferior temporal gyrus; *FuG*, fusiform gyrus; *PhG*, hippocampal gyrus; *pSTS*, posterior superior temporal sulcus; *IPL*, inferior parietal lobule. *PoG*, postcentral gyrus. The ID is the number of the parcel in the Brainnetome atlas (Fan et al., 2016). Here we only summarized the language-related behavioral domains and paradigm classes; the full behavioral domains and paradigm classes for each node are available at <http://atlas.brainnetome.org/bnatlas.php>.

Supplementary Table 2. Loading weight of each task on each component in principle component analysis.

| **Tasks** | **General Semantics** | **Elementary Perception** | **Phonological Processing** |
| --- | --- | --- | --- |
| Word-picture verification | 0.922 | 0.28 | -0.024 |
| Word associative matching | 0.837 | 0.271 | -0.152 |
| Picture associative matching | 0.675 | 0.377 | 0.242 |
| Oral picture naming | 0.932 | 0 | 0.071 |
| Naming to definition | 0.873 | -0.197 | 0.232 |
| Oral sound naming | 0.712 | -0.251 | 0.416 |
| Visual perception | 0.074 | 0.8 | -0.059 |
| Sound perception | 0.036 | 0.72 | 0.158 |
| Auditory lexical decision | 0.091 | 0.138 | 0.943 |
